# Spatially resolved epigenomic profiling of single cells in complex tissues

**DOI:** 10.1101/2022.02.17.480825

**Authors:** Tian Lu, Cheen Euong Ang, Xiaowei Zhuang

**Affiliations:** Howard Hughes Medical Institute, Harvard University, Cambridge, MA 02138, USA; Department of Chemistry and Chemical Biology, Harvard University, Cambridge, MA 02138, USA; Department of Physics, Harvard University, Cambridge, MA 02138, USA

## Abstract

The recent development of spatial omics methods enables single-cell profiling of the transcriptome and the 3D genome organization in a spatially resolved manner. Expanding the repertoire of spatial omics tools, a spatial epigenomics method will accelerate our understanding of the spatial regulation of cell and tissue functions. Here, we report a method for spatially resolved profiling of epigenomes in single cells using in-situ tagmentation and transcription followed by highly multiplexed imaging. We profiled histone modifications marking active promoters and enhancers, H3K4me3 and H3K27ac, and generated high-resolution spatial atlas of hundreds of active promoters and putative enhancers in embryonic and adult mouse brains. Our results further revealed putative promoter-enhancer pairs and enhancer hubs regulating the expression of developmentally important genes. We envision this approach will be generally applicable to spatial profiling of epigenetic modifications and DNA-binding proteins, advancing our understanding of how gene expression is spatiotemporally regulated by the epigenome.

## INTRODUCTION

Spatiotemporal control of gene expression is essential for the development and function of cells and tissues. The regulatory information encoded in the epigenome, such as histone and DNA modifications, enables the same set of genes situated in the genome to be differentially activated or repressed to generate different types of cells during development; and mis-regulation of gene expression leads to diseases (Allis and Jenuwein, 2016; Henikoff and Smith, 2015; Moris et al., 2016; Zhao et al., 2021; Zoghbi and Beaudet, 2016). Sequencing-based approaches have been traditionally used to profile histone and DNA modifications in a high-throughput manner in an ensemble of cells. Recently, epigenetic sequencing techniques have been extended to the single-cell level to enable the characterizations of chromatin accessibility and epigenetic modifications in individual cells (Bartlett et al., 2021; Bartosovic et al., 2021; Buenrostro et al., 2015; Carter et al., 2019; Cusanovich et al., 2015; Gravina et al., 2016; Kaya-Okur et al., 2019; Wang et al., 2019b; Wu et al., 2021; Zhu et al., 2021)

However, the spatial context of cells, which are critical for understanding tissue development and function, are lost in sequencing-based methods that require cell dissociation. Spatially resolved single-cell profiling of epigenetic properties, such as epigenetic modifications marking active enhancers and promoters, will greatly facilitate our understandings of how the epigenome shapes the development of cell types and control of cell states in the native context of complex tissues. For example, during embryonic brain development, morphogenic gradients and transcription factors form complex spatial patterns, giving rise to a myriad of neural progenitors destined to become different types of neurons and non-neuronal cells (Cadwell et al., 2019; Gelman et al., 2012; Hébert and Fishell, 2008; Molnár et al., 2019; O’Leary et al., 2013; Rakic, 2009). Recent evidence suggests that diverse enhancer recruitments may help generate finely delineated domains or protodomains within the developing brain, fine-tuning the broad patterns generated by transcription factors and morphogenic gradients (Pattabiraman et al., 2014; Visel et al., 2013). Progenitors from those finely delineated domains have been shown to give rise to different neuronal subtypes in various brain regions, highlighting the need for epigenomic mapping with high spatial resolution (Silberberg et al., 2016). Moreover, in the adult brain, neurons from different subtypes and cortical layers were found to have different chromatin accessibilities and epigenetic modification profiles (Gray et al., 2017; Graybuck et al., 2021; Mo et al., 2015; Zhu et al., 2021), shedding light on how spatially varying epigenetic properties regulate gene expression. Tens to hundreds of thousands of epigenetic elements, such as putative active enhancers, have been identified in both embryonic and adult brains (Gorkin et al., 2020; Gray et al., 2017; Graybuck et al., 2021; Mo et al., 2015; Preissl et al., 2018; Shen et al., 2012; Visel et al., 2009, 2013; Yue et al., 2014), but the detailed spatial distributions remain unclear for most of these elements. High-throughput and high-resolution spatial profiling of these epigenetic elements will greatly facilitate the functional understanding of the epigenome.

The transgenic approach that delivers the enhancer sequence fused to a reporter expression cassette into the animal has been used to measure the spatial patterns of thousands of putative enhancers in the embryonic mouse brain in a heroic effort that spanned more than 10 years (Pattabiraman et al., 2014; Silberberg et al., 2016; Visel et al., 2007, 2013). This approach requires extensive cloning and generation of transgenic animals. In addition, mapping putative enhancers in a setting where the enhancer activity is shown by the adjacent reporter might not always recapitulate the endogenous epigenetic activities. A spatial profiling approach that can map the endogenous epigenetic activities, such as active enhancers and promoters, of individual cells in a high throughput manner is thus highly desirable. Moreover, it is essential that such a spatial epigenomics approach has a high genomic resolution because these epigenetic elements are typically short (∼1 kb or shorter).

Recently, spatial genomics approaches have been developed to profile the transcriptome using either imaging-based approaches (multiplexed FISH or *in situ* sequencing) with single cell resolution (Lein et al., 2017; Zhuang, 2021) or spatially resolved RNA capture following by sequencing (Larsson et al., 2021). The imaging-based approaches have also allowed the 3D organization of the DNA in single cells to be measured at the genome scale, imaging thousands of chromatin loci with a genomic resolution on the order of tens of kilobases to megabases (Payne et al., 2021; Su et al., 2020; Takei et al., 2021a, 2021b). Such genome-scale chromatin imaging has also been combined with protein imaging to study the spatial relationship between chromatin loci and nuclear structures, including various nuclear bodies and histone marks (Su et al., 2020; Takei et al., 2021a, 2021b), but due to limited imaging resolution, it is difficult to determine whether chromatin loci that colocalize with the histone marks carry these marks or are just in spatial proximity to them. Expansion microscopy can improve the accuracy in determining the epigenetic state of chromatin loci and has been demonstrated for imaging the histone modifications of a few genomic loci at 10-kb resolution (Woodworth et al., 2021). However, a technique that allows epigenetic-state imaging of the chromatin in individual cells with high genomic resolution and high genomic throughput is still in demand.

Here, we developed an imaging method to measure the epigenetic modifications of chromatin in individual cells in a spatially resolved manner with high genomic throughput and high genomic resolution. We demonstrated the ability to image genomic loci as short as a few hundred bases, identifying their epigenetic states and mapping their spatial distributions in tissues. We used this approach to map hundreds of active promoters and putative enhancers marked by specific histone modifications in mouse embryonic and adult brains. Our imaging data not only confirmed previously known spatial patterns of promoters and enhancers in the brain, which validates our method, but also revealed previously unknown high-resolution spatial distributions of putative enhancers and predicted novel enhancer-promoter pairs and enhancer hubs for regulating developmentally important genes in the embryonic brain.

## RESULTS

### Epigenomic MERFISH enables *in situ* spatially resolved single-cell profiling of epigenetic modifications

In order to image chromatin loci with specific epigenetic modifications in a high throughput manner, we first captured the epigenetic modifications on the chromatin *in situ* and tagged the DNA with T7 promoters at or near the modification sites (**Figure 1A**). We then performed *in situ* transcription of the T7 tagged regions to generate RNAs *in situ* (**Figure 1A**). Finally, we detect the transcribed RNAs by multiplexed error robust fluorescence *in situ* hybridization (MERFISH) (**Figure 1A**), an imaging method that allows RNA imaging at the transcriptomic scale (Chen et al., 2015), which in turn allowed us to image a large number of epigenetically modified chromatin loci simultaneously. Hereafter, we referred to this method as epigenomic MERFISH.

**Figure 1.**
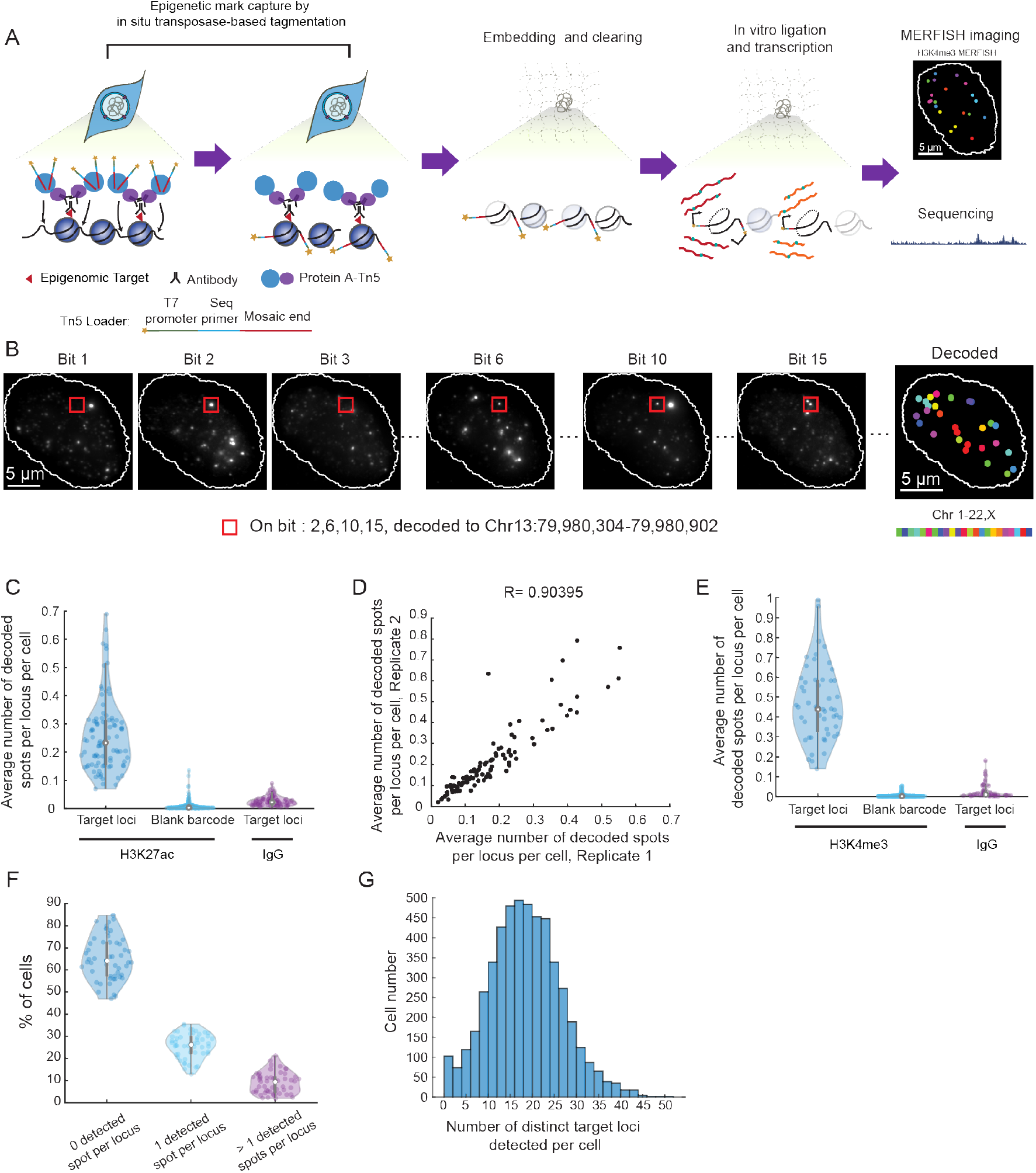
Spatially resolved single-cell profiling of epigenetic modifications by epigenomic MERFISH. (A) Schematic showing the workflow of epigenomic MERFISH. Cells were fixed, permeabilized, and treated with primary antibodies recognizing the epigenetic modifications of interest, secondary antibodies, and protein A coupled transposase (PA-Tn5) to generate DNA fragments tagged with T7 promoter and sequencing primers. The sample was embedded into polyacrylamide gel and cleared, while the tagged DNA are crosslinked to the gel via the acrydite group in the tag. The tagged DNA fragments were then transcribed into RNAs by T7 polymerase. The resulting RNA was either detected by MERFISH imaging or were reversed transcribed and subjected to sequencing. (B) Epigenomic MERFISH image of 90 target H3K27ac loci in a single cell. The images from individual bits are shown on the left. The decoded image is shown on the right, with individual spots color-coded based on the chromosomal identities of the genomic loci. The spot, marked by the red box, was observed in bit 2, 6, 10, 15 and hence decoded to the Chr13: 79,980,304-79,980,902 locus. (C) Violin plot showing the average number of decoded spots per cell for each target H3K27ac locus (left) and each blank barcode (middle) when H3K27ac antibody is used to capture the epigenetic mark. Also shown is the violin plot of the average number of decoded spots per cell for each target H3K27ac locus when a control IgG is used instead the H3K27ac antibody (right). Each dot in the violin plots correspond to a single H3K27ac locus or a blank barcode. (D) Scatter plot showing the correlation between two biological replicates of H3K27ac imaging. Each dot corresponds to a single H3K27ac locus. (E) Same as (C) but for epigenomic MERFISH imaging of 52 target H3K4me3 loci that correspond to the promoters of 52 essential genes. (F) Violin plot showing the percentage of cells with 0, 1, or >1 detected spots for individual target H3K4me3 loci. The target loci correspond to the promoters of 52 essential genes. Each dot in the violin plots corresponds to a single H3K4me3 locus. For a given locus, ∼36% (median percentage across 52 loci) of cells showed at least one detected spot. (G) Histogram of the number of distinct target H3K4me3 loci detected per cell. The median number of distinct target loci detected per cell is 18, which is ∼35% of the 52 total target loci.

We first optimized and validated this method in cultured hTERT-RPE1 cells. To capture the epigenetic modifications, we fixed the samples and labeled the epigenetic modification of interest using antibodies that recognize the modification (**Figure 1A**). The antibodies where then bound by protein A fused with transposase Tn5 (PA-Tn5), which allowed Tn5 to transpose the T7 promoters into the DNA region at or near the epigenetic modification site (**Figure 1A**). This procedure resulted in fragments of chromatin encompassing the epigenetic loci of interest tagged by the T7 promoters and sequencing primers at both ends, where primer tags allowed the PCR amplification and sequencing of DNA fragments, as previously done in the CUT&Tag approach (Bartlett et al., 2021; Bartosovic et al., 2021; Harada et al., 2019; Kaya-Okur et al., 2019; Liu et al., 2020; Wang et al., 2019b; Wu et al., 2021; Zhu et al., 2021). The T7 promoter tags further allowed the DNA fragments to be transcribed into RNA by the T7 polymerase for *in situ* amplification and detection. Tagmentation with the T7 promoter has also been used for signal amplification in CUT&Tag recently (Bartlett et al., 2021).

To ensure efficient and faithful capture of the epigenetic modifications in fixed cells, we screened fixation conditions using various crosslinking or precipitating fixatives and fixation durations and found that light PFA fixations with HCl treatment enabled accurate transposition near the target epigenetic loci (**Figure S1A and S1B**).

After the DNA fragments were generated by Tn5 transposition, those tagged fragments were transcribed into RNAs using the T7 RNA polymerase. This *in situ* transcription step amplifies a single copy of DNA fragment into many copies of RNA, which not only greatly increases the signal of epigenomic loci to confer detection specificity, but also allows us to detect short DNA locus that would otherwise be difficult to image by FISH. To ensure efficient transcription, we embedded the sample in polyacrylamide gel and digested the sample by proteinase K in order to remove DNA-interacting proteins that could impede T7 transcription (**Figure 1A**). This embedding and clearing procedure improved T7 amplification and generated more RNAs for better capturing of the histone modification peaks (**Figure S1B and S1C)**.

Finally, we used MERFISH to image the transcribed RNAs in the gel-embedded samples a highly multiplexed manner (Chen et al., 2015; Moffitt et al., 2016b, 2016a). As in our previous MERFISH measurements, we used N-bit barcodes with Hamming Distance 4 (HD 4) and Hamming weight 4 (HW 4) to allow error correction, where the length the barcodes (N) was chosen based on the number of target epigenetic loci. To avoid crowdedness of the FISH signal in each bit such that individual loci could be clearly resolved, we assigned barcodes to the target epigenetic loci in a manner such that only 3-5 loci in each chromosome was imaged in each bit (See STAR Methods for details in MERFISH probe design, Table S1 for oligonucleotide sequences, and Table S2 for barcode assignments).

Before performing the *in situ* MERFISH imaging of the target loci, we first collected and measured the transcribed RNAs in an untargeted manner by sequencing to test whether the epigenetic loci were faithfully captured. To this end, we measured the profiles of two histone modifications, H3K27ac and H3K4me3, in hTERT-RPE1 cells. H3K4me3 is known to be a canonical marker for active promoters; H3K27ac can mark both active promoters and enhancers and intergenic H3K27ac loci are often used to predict putative active enhancers. We found that the length distribution of the RNAs generated by *in situ* Tn5 transposition and T7 transcription was around 100-1000 bases (**Figure S1D**). The RNAs were subsequently reverse transcribed, PCR-amplified and sequenced. The genome-wide profiles of H3K27ac and H3K4me3 measured using this *in situ* tagmentation and transcription approach agreed with the H3K27ac and H3K4me3 peaks detected by ChIP-Seq and CUT&Tag, albeit with a lower peak height (**Figure S1B and S1C**).

Next, we performed MERFISH imaging of the transcribed RNAs *in situ* to achieve spatial profiling of the epigenetic loci of interest. To this end, we first selected 90 H3K27ac-positive loci in Chr1-22 and ChrX of hTERT-RPE1 cells based on the H3K27ac peaks in ChIP-seq data and designed MERFISH probes targeting these loci (**Table S1**). The selected H3K27ac loci had a median length of 500 bp. We measured these loci with a 24-bit, HW 4 and HD 4 code (**Table S2**) using 8 rounds of 3-color imaging. Among the 366 total valid barcodes, 90 were assigned to the target H3K27ac loci and the remaining ones were unassigned (referred to as blank barcodes), which allowed us to assess the mis-identification rate. The MERFISH images showed clear and decodable spots (**Figure 1B**). The spots that were decoded into individual target H3K27ac loci were on average ∼30 fold more abundant than the spots that were decoded into individual blank barcodes (**Figure 1C**), indicating a low misidentification rate. As a further control, we replaced the H3K27ac antibody with a control IgG antibody and observed a ∼10-fold reduction in the number of spots decoded into the target loci, indicating that our epigenomic MERFISH measurement results were specific to H3K27ac modifications (**Figure 1C**). Furthermore, the results were reproducible between biological replicates (**Figure 1D**).

We next estimated the detection efficiency of our epigenomic MERFISH measurements by imaging the promoters of essential genes, which should in principle be active and hence H3K4me3-positive in every cell. If the detection efficiency of epigenomic MERFISH was 100%, we should detect at least one H3K4me3-positive spot for each promoter in every cell. We selected 52 such essential genes and designed probes targeting the ±2 kb region of their transcription starting site (TSS), which presumably covers the promoter regions of the genes (**Tables S1 and S2**). Again, comparisons with the blank barcode counts and with the measurements using the control IgG showed that our detection of the H3K4me3-positive loci was highly specific with a low mis-identification rate (**Figure 1E**). For any given target locus, ∼36% (median across 52 target loci) of the cells exhibited at least one detected spot (**Figure 1F**), and in any given cell, ∼35% (median across ∼5000 cells) of the target loci were detected (**Figure 1G**). This detection efficiency was higher than the detection efficiencies of single-cell CUT&Tag (∼1-8%) (Bartosovic et al., 2021) and single-cell ATAC-seq (∼5-10%) (Chen et al., 2019; Fang et al., 2021).

### Region- or layer-specific patterns of active promoters in the mouse brain

Next, we demonstrated the spatial profiling power of epigenomic MERFISH by using it to map histone modifications in mouse brain tissues. As a proof of principle, we first imaged active promoters marked by H3K4me3 for genes with known spatial expression patterns in the embryonic and/or adult brain. We targeted the H3K4me3 peaks near the TSS of 127 genes, including genes that exhibit region-specific expression in the embryonic brain (*Foxg1, Emx2, Gbx2, En2, etc*), genes that exhibit layer-specific expression in the adult cortex (*Cux2, Rorb, Fezf2, etc*), and some well-known neuron subtype markers (*Slc17a7, Gad1, Gad2, etc*) (**Tables S1 and S2**).

Upon further optimization of the epigenomic MERFISH protocol for tissue slices (see STAR Methods), we mapped the spatial distributions of these H3K4me3-positive loci in brain tissue slices of adult (**Figure 2A**) and embryonic day 13.5 (E13.5) (**Figure 3A**) mice. As in the experiments for cultured cells, we observed clear and decodable spot in individual cells in the mouse brain tissue (**Figure S2A**), and the detection of the H3K4me3-positive loci in brain tissues were highly specific to the H3K4me3 antibody with a low mis-identification rate (**Figure S2B**) and reproducible between replicates (**Figure S2C**).

**Figure 2.**
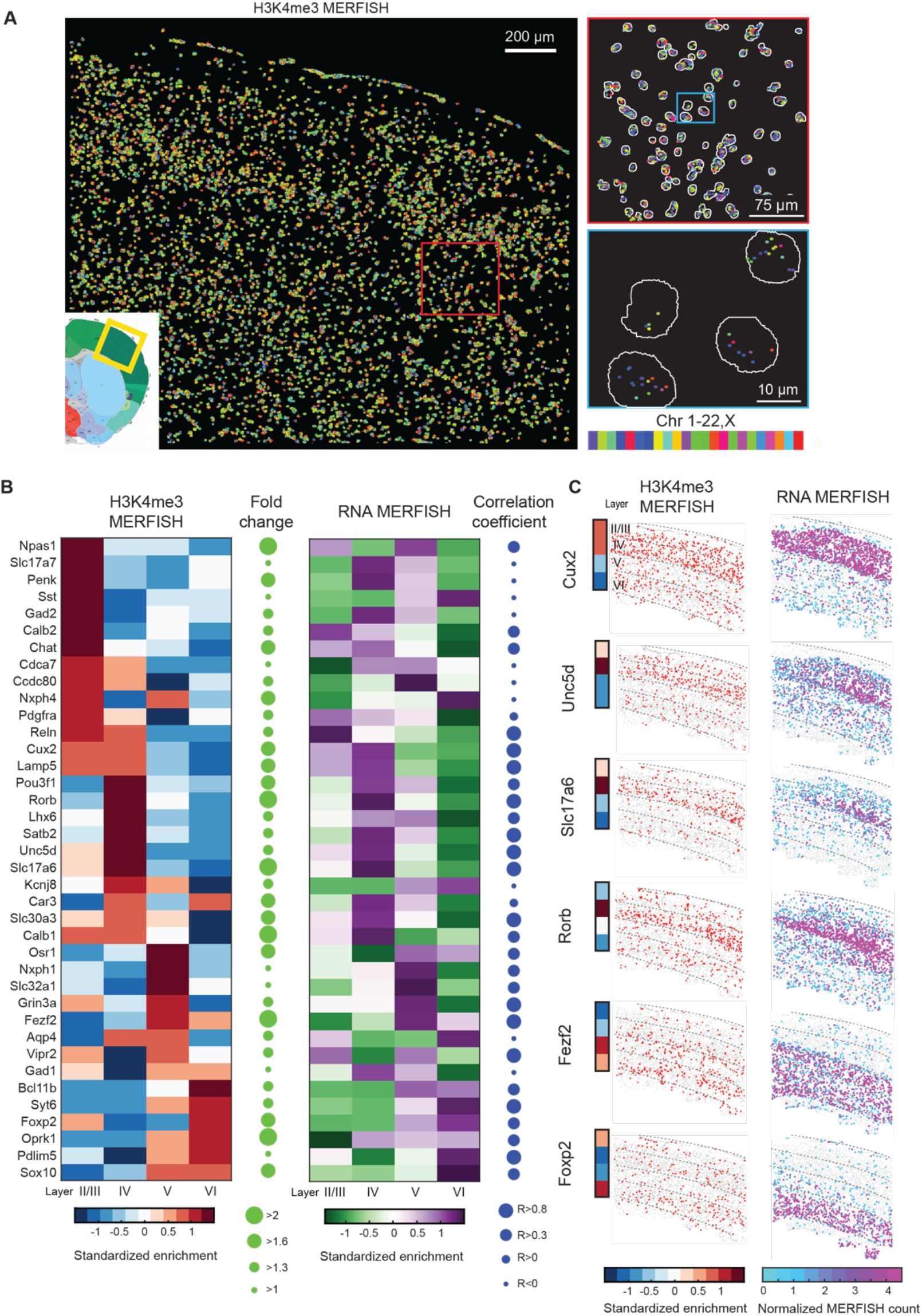
Spatially resolved single-cell profiling of layer-specific active promoters in adult mouse cortex. (A) Left: Epigenomic MERFISH image of 127 target H3K4me3 loci in the somatosensory cortex region of a coronal slice of an adult mouse brain. Top right: A magnified view of the red-boxed region from the left panel showing the decoded spots of H3K4me3 loci in individual cells. Bottom right: A magnified view of the blue-boxed region from the top right panel. Segmentation of individual nuclei are shown in white and decoded spots are color-coded by the chromosomal identities of the genomic loci. (B) Left: Heatmap of the standardized layer enrichment for the promoter H3K4me3 signals measured by epigenomic MERFISH for the indicated genes. A cell is considered H3K4me3-positive for the promoter locus of a gene if at least one decoded spot for this locus is detected in the cell. For each promoter locus, the standardized layer enrichment in a specific layer is calculated as the z-score of the following quantity: the fraction of cells in the layer that is H3K4me3-positive for this locus, and is presented in colors based on the colored scale bar at the bottom. Middle Left: Fold change in the promoter H3K4me3 signals between the layers with the maximum and minimum enrichment. The size of the circle reflects the value of the fold change. Middle Right: Heatmap of the standardized layer enrichment for the RNA expression level measured by RNA MERFISH for the indicated genes. For each gene, the standardized layer enrichment in a specific layer is calculated as z-score of the following quantity: the fraction of cells in the layer that express this gene, and is presented in color based on the colored scale bar shown at the bottom. Right: Correlation of layer enrichment between the epigenomic MERFISH and RNA MERFISH data. The size of the circle reflects the value of the Pearson correlation coefficient R. (C) Left: Epigenomic MERFISH images showing layer enrichment of H3K4me3 signals for the promoters of six indicated genes. Each dot in the images represent a cell and red dots represent cells with positive H3K4me3 signals. The layer enrichment heatmap on the left is presented in colors according to the colored scale bar shown at the bottom and is reproduced from panel (B). Right: RNA MERFISH images showing layer enrichment of RNA expression for the six indicated genes (Zhang et al., 2021). The RNA expression level in each cell is presented in colors according to the colored scale bar shown at the bottom.

**Figure 3.**
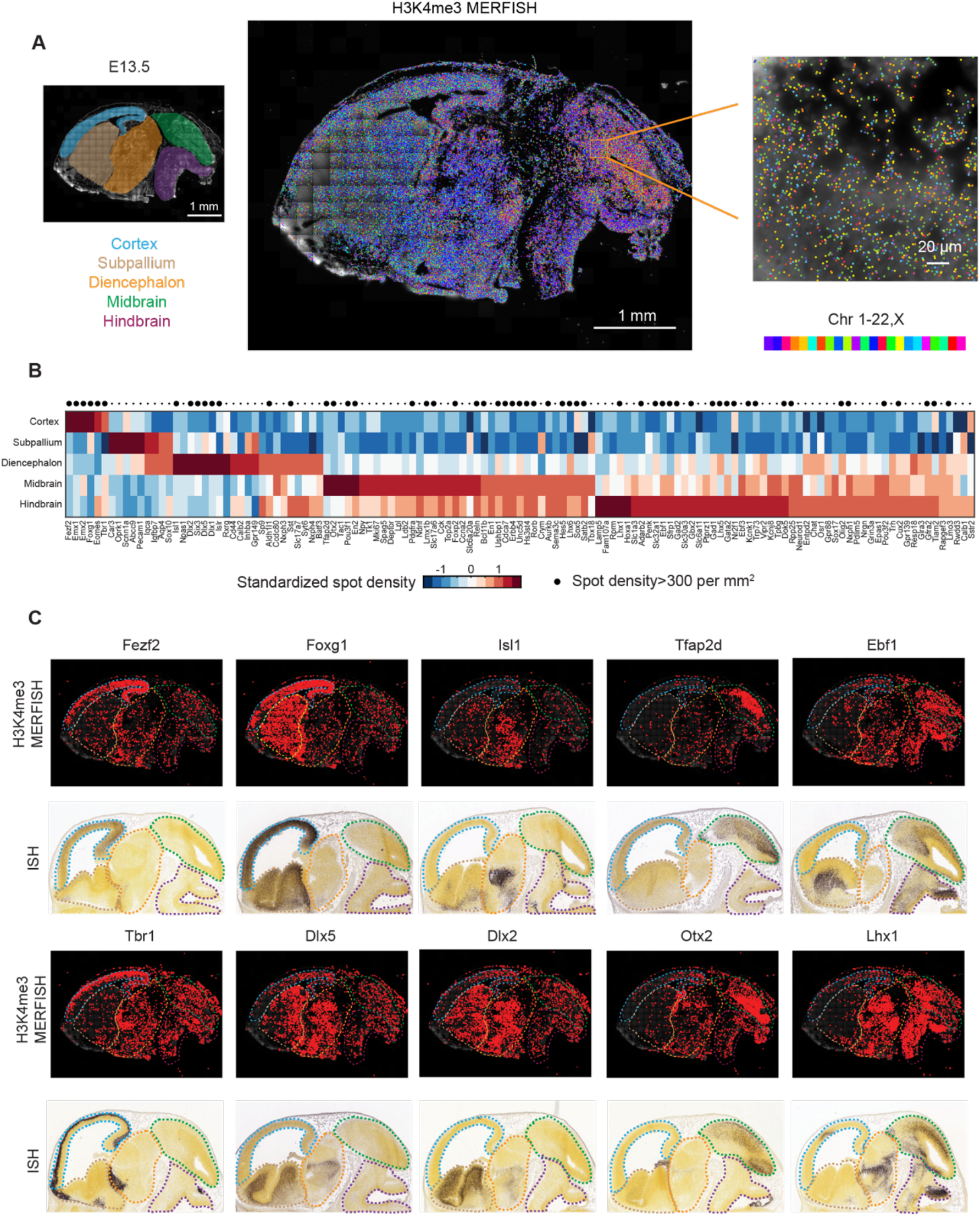
Spatially resolved profiling of active promoters in mouse embryonic brain. (A) Left: Schematic highlighting different brain regions, cortex, subpallium, diencephalon, midbrain, hindbrain, of an imaged sagittal slice of a E13.5 mouse brain. The background shows the DAPI signal. Middle. Epigenomic MERFISH image of 127 target H3K4me3 loci in the slice. Right: An enlarged region for orange box in the midbrain. All decoded spots are plotted on the background of the DAPI signal and are color-coded by the chromosomal identities of the genomic loci. (B) Heatmap showing the standardized region enrichment for each of the 127 target H3K4me3 loci in different brain regions. The brain region is segmented manually based on the cytological hallmarks. The spot density in each region is measured by the number of decoded spots in this region divided by the DAPI-positive area. The number was then standardized for each locus across the five brain regions such that the mean of the enrichment for each locus is 0 and the variance is 1 to give the standardized region enrichment, which is shown in color based on the colored scale bar at the bottom. Top: Large dot indicating the loci that have a H3K4me3 spot density larger than 300 per mm^2^. (C) Epigenomic MERFISH images of the H3K4me3 signals of the promoters of ten transcription factors shown in comparison with the Allen Brain ISH images showing the expression patterns of the corresponding genes.

We then characterized the spatial distributions of the H3K4me3 signals of these 127 loci in adult mouse cortex, focusing on whether loci corresponding to the promoters of cortical-layer marker genes exhibited the expected layer-specific enrichment of H3K4me3 signals. We imaged these 127 loci in thousands (∼4,000) of individual cells in the somatosensory cortex of adult mouses brains (**Figure 2A**), and compared the spatial patterns of the detected H3K4me3 loci with the expression pattern of their corresponding genes that we recently measured by MERFISH (Zhang et al., 2021). Of the 127 H3K4me3 loci probed here, 38 loci had corresponding genes in the published RNA MERFISH data, and the layer enrichment pattern that we observed for these loci and genes were largely similar between the epigenomic MERFISH and RNA MERFISH measurements (**Figure 2B**). For example, like RNA MERFISH signals, *Cux2 and Unc5d* promoter H3K4me3 signals measured by epigenomic MERFISH were enriched in layers II/III and IV, *Rorb* and *Slc17a6* promoter H3K4me3 signals were enriched in layer IV, *Fezf2* promoter H3K4me3 signal was enriched in layers V and VI, and *Foxp2* promoter H3K4me3 signal was enriched in layer VI (**Figure 2C**).

We noticed that for some loci, the H3K4me3 signals showed apparently different distributions as compared to the distributions of the corresponding genes measured by RNA MERFISH. However, the degree of enrichment for most of these H3K4me3 loci were relatively small (**Figure 2B)** and visual inspection of H3K4me3 images of these loci did not show obvious layer-specific enrichment (see examples in **Figure S3**).

Next, we characterized how the H3K4me3 signals of the 127 target loci were distributed in the embryonic brain. To identify genes with region-specific expression, we determined the detected spot density of these 127 loci in each of the five brain regions: the cortex, subpallium, diencephalon, midbrain, and hindbrain (**Figure 3A**). Many loci show enrichment of expression in specific brain regions (**Figure 3B**). To validate our results, we focused on those loci with reasonably high spot density (>300 spots per mm^2^) and compared their spatial distribution with the spatial expression patterns of the corresponding genes reported in Allen brain *in situ* hybridization (ISH) atlas. Of the 57 loci that satisfied this criterion, 46 have ISH data of corresponding genes measured in the E13.5 brain and ∼87% of them (40 loci out of 46) showed spatial distributions of H3K4me3 signals that were similar to the RNA distributions reported in the Allen Brain ISH atlas (see examples in **Figure 3C**). For example, the promoters of *Tbr1* and *Fezf2* exhibited H3K4me3 signal enrichment within the cortex, comparable to the expression patterns of *Tbr1* and *Fezf2* in the Allen ISH images (**Figure 3C**). Canonical transcription factors for cortical development (*Emx1*, *Emx2*, *Bcl11b* and *Eomes*) also showed H3K4me3 signal enrichment in the cortex (**Figure S4**). As a telencephalon marker, *Foxg1* showed expected H3K4me3 signal enrichment in the cortex and subpallium (**Figure 3C**). Several distal less homeodomain transcription factors (*Dlx1*, *Dlx2* and *Dlx5*) showed expected H3K4me3 signal enrichment in subpallium and diencephalon, in agreement with expression pattern of the genes in the Allen ISH images (**Figure 3C; Figure S4**) and consistent with the knowledge that these genes are important for forebrain inhibitory neuron development (Eisenstat et al., 1999). Like the *Dlx* genes, several canonical inhibitory neuronal markers (e.g. *Slc32a1* and *Gad2*) also showed H3K4me3 signals in the subpallium, a region that is predominantly populated by inhibitory neurons, whereas H3K4me3 signals for excitatory neuronal markers (e.g. *Slc17a6*) was expectedly lacking in this region (**Figure S4**). The promoter of *Isl1* showed H3K4me3 signal enrichment in the diencephalon, consistent with both the expression pattern of the gene shown in the Allen ISH image and the known expression of this gene in a subpopulation of differentiating hypothalamic neurons (Lee et al., 2016). Finally, the promoters of midbrain and/or hindbrain specific transcription factors (e.g.: *Tfap2d, Otx2, Ebf1*, *Lhx1*, *Erbb4* and *En2*) (Cepeda-Nieto et al., 2005; Joyner, 1996; Rhinn et al., 1998; Wang et al., 1997) showed expected H3K4me3 signal enrichment in midbrain and/or hindbrain (**Figure 3C and Figure S4**)

Overall, the agreement between our spatial profiling results of the H3K4me3-marked active promoters and the previously measured expression patterns of the corresponding genes further validated our epigenomic MERFISH measurements.

### Layer-enrichment patterns of putative active enhancers in mouse adult cortex

Next, we applied epigenomic MERFISH to spatially map the putative active enhancers in the brain. We first asked whether we could reveal the layer-specific enhancers by targeting genomic loci with the H3K27ac modification. When situated in regions away from the promoter sites, H3K27ac is often used as a marker for predicting putative active enhancers (Creyghton et al., 2010; Heintzman et al., 2009; Rada-Iglesias et al., 2011). However, layer-specific bulk H3K27ac sequencing data is not readily available. Recently, layer-specific chromatin accessibility has been profiled by ATAC-seq using FACS sorted layer-specific excitatory neurons labeled with fluorescent reporter driven by layer-specific promoters (Gray et al., 2017; Graybuck et al., 2021). We thus used the ATAC-seq data to guide our selection of target genomic loci, as ATAC-seq peaks that do not correspond to gene promoters are often considered possible candidates for enhancers. We selected 139 ATAC peaks that are >2kb away from known TSS of genes, show signal enrichment in one cortical layer and have non-zero reads from bulk H3K27ac ChIP-Seq data. The median length of these peak is ∼250 bp.

We performed epigenomic MERFISH imaging of these loci, targeting the H3K27ac modification, in adult mouse coronal sections containing the somatosensory cortex and profiled ∼3,600 individual cells in this region (**Figure 4A**). The results were specific to the H3K27ac antibody and consistent between replicates (**Figure S5A and S5B**). Among 139 target loci, 35 of them showed a statistically significant layer-specific pattern (**Figure 4B**). The observation that many of the loci did not show layer-specific enrichment of H3K27ac signals is not surprising considering that a substantial fraction (∼50%) of the ATAC-seq peaks are not overlapping with H3K27ac peaks measured by ChIP-seq (Fulco et al., 2019; Gray et al., 2017). Hence, the signal of ATAC peaks might not always reflect the H3K27ac level of those regions. Indeed, the 35 loci that exhibited significant layer-specific enrichment had a higher average H3K27ac signal than those not exhibiting significant layer-specific enrichment (**Figure S6**). Among these 35 loci, the layer-enrichment patterns were largely similar to those obtained from ATAC-seq (**Figure 4B**).

**Figure 4.**
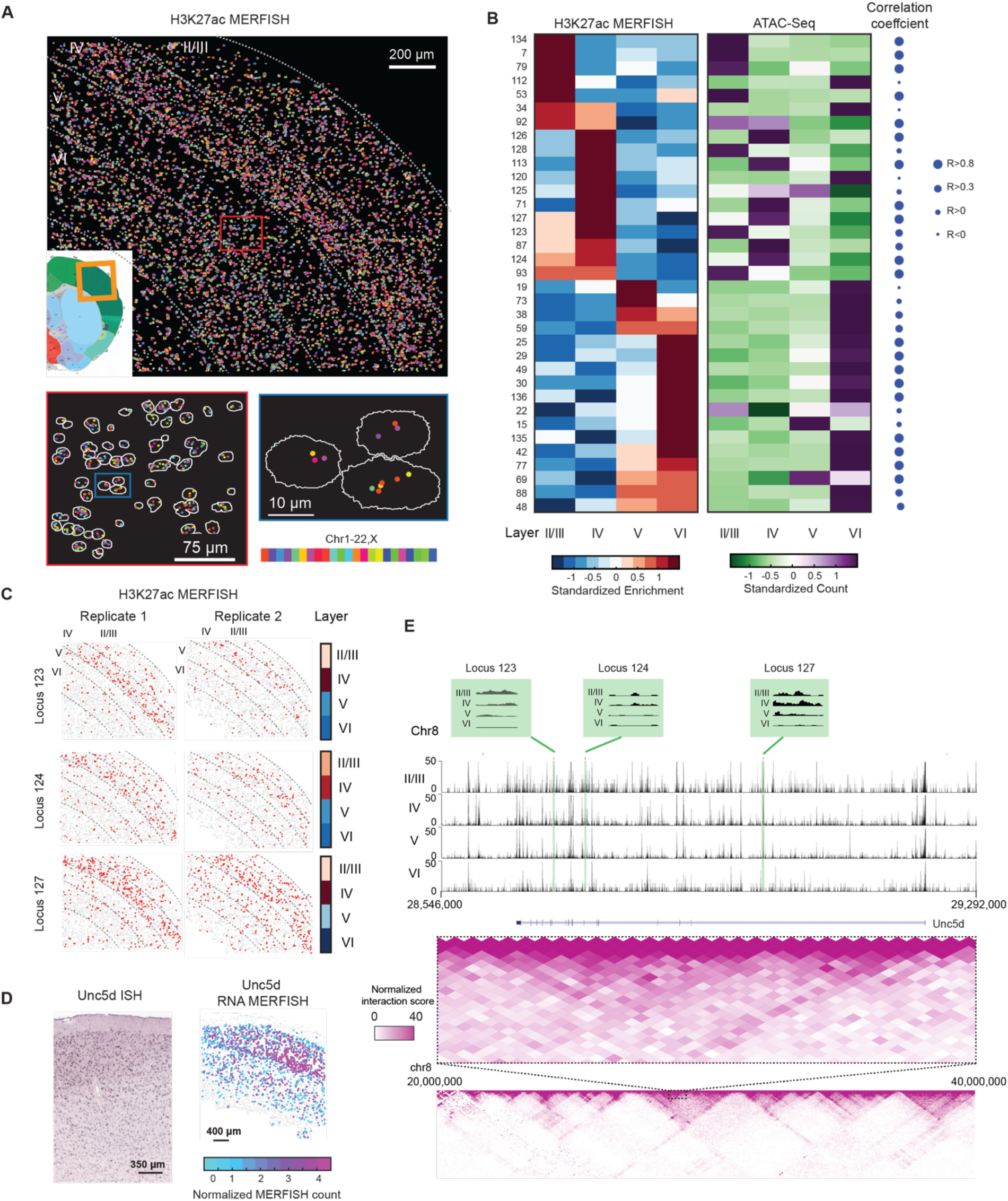
Spatially resolved single-cell profiling of layer-specific putative active enhancers in adult mouse cortex. (A) Top: Epigenomic MERFISH image of 139 target H3K27ac loci in the somatosensory cortex region of a coronal slice of an adult mouse brain. Bottom left: A magnified view of the red-boxed region from the top panel showing the decoded spots of H3K27ac loci in individual cells. Bottom right: A magnified view of the blue-boxed region from the top right panel. Segmentation of individual nuclei are shown in white and decoded spots are color-coded by the chromosomal identities of the genomic loci. (B) Left: Heatmap showing the standardized layer enrichment for the H3K27ac signal measured by epigenomic MERFISH for the indicated genomic loci. Standardized layer enrichment is calculated as described in Figure 2B and shown in color based on the colored scale bar at the bottom. Middle: Heatmap of the corresponding standardized reads per million for each of target loci from published layer specific ATAC-seq data (Gray et al., 2017), shown in color based on the colored scale bar at the bottom. Right: Correlation between the layer enrichment derived from epigenomic MERFISH data and ATAC seq data. The size of the circle reflects the value of the Pearson correlation coefficient, R. (C) Epigenomic MERFISH images of the H3K27ac signals for the three target loci showing their enrichment in layers II/III and IV. Images of two replicates are shown. Quantification of the layer enrichment are shown on the right (reproduced from Figure 4B). (D) ISH and RNA MERFISH images of the RNA expression level of the *Unc5d* gene. In the RNA MERFISH image (Zhang et al., 2021), the RNA expression level in each cell is color coded according to the colored scale bar shown at the bottom. (E) Top: UCSC browser track of the ATAC-seq data (Chr8: 28,600,000 - 29,250,000) (Gray et al., 2017) showing the location of the three target loci (loci 123, 124, 127) in the intronic regions of Unc5d, which are 520,651 bp, 476,571 bp, and 226,831 bp downstream of *Unc5d* TSS. Regions marked in green are the three target loci with the green boxes above showing the enlarged version of the ATAC-seq track of the marked loci. Bottom: Hi-C map of a genomic region (Chr8: 20,000,000 – 40,000,000) harboring the Unc5d loci (50-kb resolution, Knight-Ruiz normalized) obtained from the mouse brain (Deng et al., 2015), with the Chr8: 28,600,000 - 29,250,000 region corresponding to a sub-TAD enlarged and shown above (25-kb resolution, Knight-Ruiz (KR) normalized).

Notably, some of these putative enhancer loci exhibited layer-specific enrichment patterns of H3K27ac signals that were similar to the spatial expression patterns of nearby genes. For example, the three putative enhancer loci (loci 123, 124 and 127) within 600 kb of the TSS of gene *Unc5d* showed a consistent and significant enrichment in layers II/III and IV (**Figure 4C**), and the *Unc5d* gene also showed enriched expression in layers II/III and IV (**Figure 4D**). The existing Hi-C data of the mouse brain (Deng et al., 2015) showed that these three loci and the *Unc5d* gene are located within the same sub-TAD (**Figure 4E**). These results suggest that loci 123, 124 and 127 are putative enhancers for the *Unc5d* gene and that the spatially profiling power of epigenomic MERFISH could help identify putative promoter-enhancer pairs.

### High-resolution spatial profiling of putative active enhancers in mouse embryonic brains

We next imaged putative active enhancers marked by H3K27ac in the E13.5 embryonic mouse brain to identify region-specific spatial patterns of enhancer activity (**Figure 5A**). To this end, we targeted a total of 142 H3K27ac-positive loci and five loci with low H3K27ac counts as negative controls, selected based on previous ChIP-Seq data obtained from embryonic brain (Gorkin et al., 2020). The five negative control loci showed a comparable number of detected spots to those of blank barcodes and ∼10-fold fewer detected spots compared to the H3K27ac-positive loci, indicating a low false positive detection rate (**Figure S5C**). As a further validation, we compared our results with previous ChIP-Seq data obtained from E13.5 forebrain, midbrain and hindbrain (Gorkin et al., 2020) by grouping the epigenomic MERFISH signals into these three major brain regions and we observed a similar region-specific enrichment pattern to the ChIP-Seq results (**Figure 5B**).

**Figure 5.**
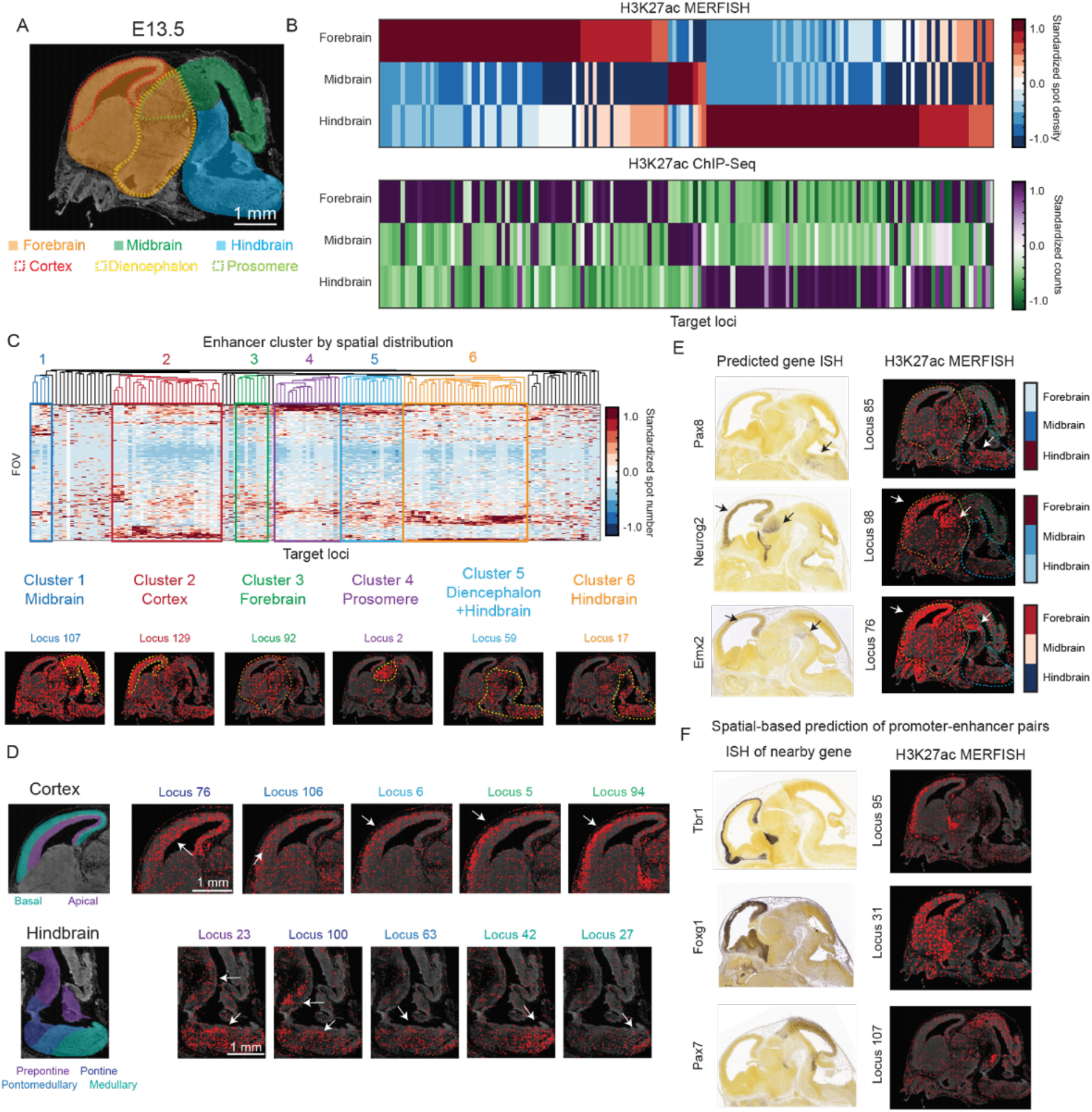
Spatially resolved profiling of putative active enhancers in mouse embryonic brain. (A) Top: Schematic highlighting different brain regions (forebrain, midbrain, and hindbrain in solid color shades and cortex, diencephalon and prosomere in dotted color lines) of an imaged sagittal slice of a E13.5 mouse brain. The background shows the DAPI signal. (B) Top: Heatmap showing the standardized region enrichment for the H3K27ac signal measured by epigenomic MERFISH for 142 target genomic loci. Standardized region enrichment is calculated as described in Figure 3B and shown in color based on the colored scale bar on the right. Bottom: Heatmap showing the corresponding standardized reads per million for the target loci from published E13.5 H3K27ac ChIP-seq data (Gorkin et al., 2020), shown in color based on the colored scale bar on the right. (C) Top: Hierarchical clustering of the 142 target genomic loci based on the measured spatial distributions of the H3K27ac signals of individual loci. The spatial distribution of each locus is presented as the number of H3K27ac spots in each imaged field-of-view (FOV: 0.04 mm^2^) for the locus with each FOV presented as a row. Six major clusters that contains >3 loci are shown, representing six different spatial patterns (enrichment in midbrain, cortex, forebrain, prosomere, diencephalon+hindbrain, and hindbrain). The brain regions are highlighted in Figure 5A. Bottom: Epigenomic MERFISH images of the H3K27ac signals of six representative loci, one for each of the six clusters. (D) Epigenomic MERFISH images of two clusters of loci that show fine spatial distribution changes within the cortex and hindbrain. White arrows point to the region of the H3K27ac signal enrichment. Schematic shown on the left depict the regions of interest. (E) Comparison between the spatial distributions of H3K27ac signals of three putative enhancer loci measured by epigenomic MERFISH (right) and the expression patterns of the corresponding predicted genes measured by ISH (left). Quantifications of the region-specific enrichment of the H3K27ac signals of the putative enhancers are shown on the right (reproduced from Figure 5B). White arrows point to the region of the H3K27ac signal enrichment, and dashed lines mark the boundary of fore, mid, and hindbrain in the epigenomic MERFISH images. Black arrows point to the region of the gene expression enrichment in the Allen ISH images. (F) Prediction of putative promoter-enhancer pairs using the H3K27ac epigenomic MERFISH data and the RNA expression pattern of the nearby genes. Epigenomic MERFISH of the H3K27ac signals of three target loci are shown on the right and Allen ISH images of the corresponding nearby genes are shown on the left.

To explore the spatial distributions of these putative active enhancers at a high spatial resolution, we performed hierarchical clustering analysis of these loci based on the spatial distributions of their H3K27ac signals. We obtained six major clusters corresponding to loci with H3K27ac signals enriched in the following six brain regions: midbrain, cortex, forebrain, prosomere, diencephalon+hindbrain, and hindbrain (**Figure 5C,** regions marked in **Figure 5A**). To understand whether each cluster of loci was potentially recognized by a specific transcription factor, we performed motif searching analysis using MEME to find transcription factor motifs enriched within those clusters of loci. Among the six clusters, we found both known motifs for specific transcription factors, including *Ascl2*, *Rfx*, *Zfp652*, *Tcf7l2*, *Sfip1* and *Sp2* motifs enriched in midbrain, cortex, forebrain, prosomere, diencephalon+hindbrain and hindbrain clusters respectively, as well as previously unknown motifs (with top two motifs shown for each cluster in **Figure S7**).

Visual inspection of the H3K27ac signals for loci within individual clusters further revealed more refined spatial patterns (**Figure 5D**). For example, a set of putative enhancers in the cortex cluster (loci 76, 106, 6, 5, 94) showed progressive changes in their spatial distributions from the apical to the basal side of the cortex with loci 76 signal distributed in the apical side, loci 5, and 94 signals distributed in the basal side, and loci 106 and 6 in between (**Figure 5D, top**). Embryonic hindbrain develops into future pons, cerebellum and medulla. Several enhancers in the hindbrain cluster (loci 23, 100, 63, 42, 27) showed interesting local enrichment of H3K27ac signals within different subregions of the hindbrain (**Figure 5D, bottom**).

For most putative enhancers, the genes that they regulate remain unknown. We posit that correlation of the enhancer activity and gene expression spatial patterns can help predict enhancer-gene pairs. Among 142 putative enhancer loci that we imaged, six of them reside within the genomic regions that have previously been predicted to be putative enhancers of genes with existing ISH data in the E13.5 brain. Interestingly, of the six loci, three of them showed spatial patterns of H3K27ac signals that matched with the expression pattern of their predicted gene targets (**Figure 5E**), providing further support for these previous predictions. In addition to supporting previously predicted enhancer-gene pairs, correlation of the spatial patterns between putative active enhancers and proximal genes could also be used to generate hypothesis of promoter-enhancer pairs. For example, several putative enhancer loci (loci 95, 31, 107) in our measurements showed spatial distributions of H3K27ac signals that matched the spatial expression patterns of their nearest genes in the genomic space (*Tbr1, Foxg1 and Pax7*, respectively) (**Figure 5F**), suggesting potential regulation of these genes by these putative enhancers.

### Putative active enhancer hubs for developmentally important genes in mouse embryonic brain

The phenomenon of multiple enhancers regulating one gene has been observed in the vertebrate system, possibly ensuring transcriptional robustness during development (Frankel et al., 2010; Osterwalder et al., 2018). Such observations have also been previously reported in the invertebrate system and are referred to as the “shadow enhancers” (Hong et al., 2008).

Interestingly, within the spatial clusters of putative enhancer loci that we observed (**Figure 5C**), we often found that multiple loci within the same cluster were located near a common gene in the genomic space and showed similar spatial patterns of H3K27ac activity to the expression pattern of the gene. For example, a set of ten putative enhancer loci (Loci 66-75) in the prosomere cluster are within ± ∼300 kb genomic distance from the promoter of *Tcf7l2*, which are all located within the same sub-TAD (**Figure 6A and 6B**). All ten enhancers showed H3K27ac signals enrichment in the prosomere region, which resembled the spatial expression pattern of *Tcf7l2* (**Figure 6A**), whereas the other gene located in the same sub-TAD (*Vti1a*) (**Figure 6B**) has a different expression pattern (enriched expression in the cortex) and are known to regulate cortical development (Sokpor et al., 2021). These results suggest the possibility that these ten loci form an enhancer hub to regulate the expression of *Tcf7l2*. Interestingly, when we performed motif search for these loci, seven of the ten loci were enriched for the *Tcf7l2* motif (loci highlighted in green in **Figure 6A,** motif shown in **Figure S7**). It is thus tempting to surmise that *Tcf7l2*, a downstream transcription factor to the developmentally important Wnt signaling, binds to its own enhancers to establish a positive feedback loop to ensure its robust expression.

**Figure 6.**
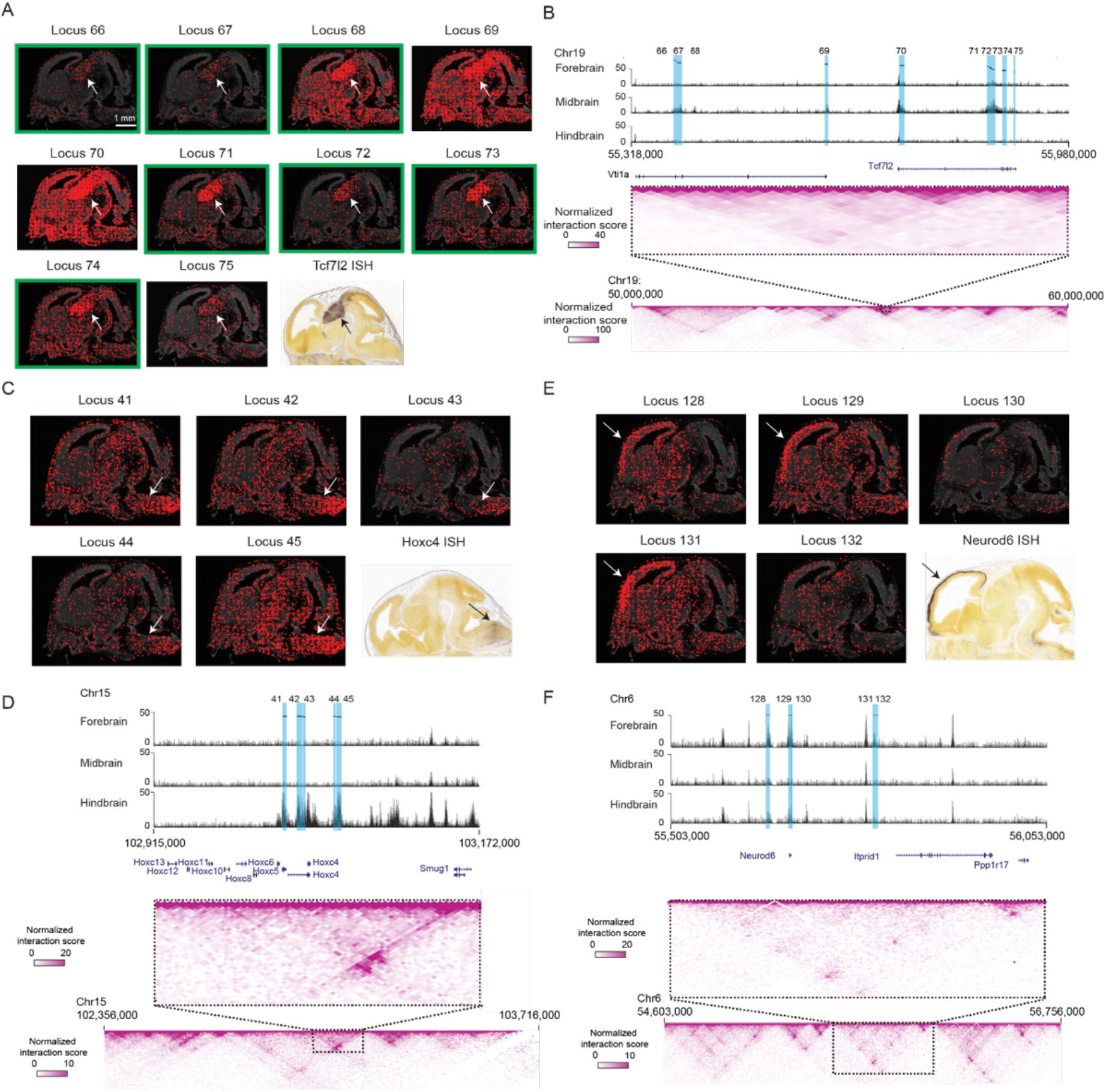
Putative active enhancer hubs for developmentally important genes in mouse embryonic brain. (A) Epigenomic MERFISH images of H3K27ac signals for 10 target loci in the prosomere cluster shown together with the Allen ISH image of the nearby gene *Tcf7l2* (bottom right). The white and black arrow points to the prosomere region where the H3K27ac signals and RNA ISH signals are most enriched, respectively. Green box marking the loci that harbors a *Tcf7l2* motif. (B) Top: H3K27ac ChIP sequencing track (Gorkin et al., 2020) of a region (Chr19:55,318,000-55,980,000) corresponding to the sub-TAD that harbors the 10 target H3K27ac loci enriched in the prosomere and the *Tcf7l2* gene. Bottom: Hi-C contact map (Deng et al., 2015) of a genomic region (Chr19:50,000,000:60,000,000) (50-kb resolution, Knight-Ruiz normalized) that includes the sub-TAD and flanking regions, with the enlarged map of the sub-TAD shown above (Chr19: 55,318,000-55,980,000) (10-kb resolution, Knight-Ruiz normalized). (C) Epigenomic MERFISH images of H3K27ac signals of 5 target loci in the hindbrain cluster shown together with the Allen ISH image of the nearby gene *Hoxc4* (bottom right). (D) Top: H3K27ac ChIP sequencing track (Gorkin et al., 2020) of a region (Chr15:102,915,000-103,172,000) corresponding to the sub-TAD that harbors the 5 target H3K27ac loci enriched in the hindbrain and several *Hoxc* genes. Bottom: Hi-C contact map (Deng et al., 2015) of a genomic region (chr15:102,356,000-103,716,000) (5-kb resolution, Knight-Ruiz normalized) that includes the sub-TAD and flanking regions, with the enlarged map of the sub-TAD shown above (Chr15:102,915,000-103,172,000) (5-kb resolution, Knight-Ruiz normalized). (E) H3K27ac epigenomic MERFISH images of 5 target loci in the cortex cluster shown together with the Allen ISH image of the nearby gene *Neurod6* (bottom right). (F) Top: H3K27ac ChIP sequencing track (Gorkin et al., 2020) of a region (Chr6:55,503,000-56,053,000) corresponding to the sub-TAD that harbors the 5 target H3K27ac loci enriched in the cortex and the *Neurod6* gene. Bottom: Hi-C contact map (Deng et al., 2015) of a genomic region (Chr6:54,603,000-56,756,000) (5-kb resolution, Knight-Ruiz normalized) that includes the sub-TAD and flanking regions, with the enlarged map of the sub-TAD shown above (Chr6:55,503,000-56,053,000) (5-kb resolution, Knight-Ruiz normalized).

Similarly, we found five putative enhancer loci in the hindbrain cluster (Loci 41-45) near the promoter of *Hoxc4*, which is known to express in the hindbrain (**Figure 6C and 6D**), and five putative enhancer loci in the cortex cluster (Loci 128-132) near the promoter of *Neurod6*, which is known to express in the cortex (**Figure 6E and 6F**). In both cases, the putative enhancer loci resided in the same sub-TAD with the genes (**Figure 6D and 6F**) and exhibited spatial patterns of H3K27ac signals that were similar to the spatial expression pattern of the gene (**Figure 6C and 6E**). Like the prosomere cluster describe above, these cortex and hindbrain clusters may also form enhancer hubs to regulate the expression of the corresponding *Hoxc4* (potentially some other *Hoxc* genes as well) and *Neurod6* genes. Together, these results suggest that spatially resolved epigenetic profiling could be used to predict putative enhancer hubs for regulating gene expression.

## Discussion

In this work, we developed a method – epigenomic MERFISH – for spatially resolved single-cell epigenomic profiling. In this method, we captured the epigenomic marks of interest *in situ* using transposase-based tagmentation of T7 promoter, amplified the tagged DNA fragments carrying the epigenomic marks using *in situ* transcription, and then detected the resulting RNA molecules using MERFISH imaging. Using this approach, we demonstrated the ability to profile the sites of epigenetic modifications on chromatin in individual cells with high spatial and genomic resolution, as well as high genomic throughput. Histone modifications on genomic loci as short as a few hundred bases can be imaged, providing a genomic resolution of <1kb. In our proof-of-principle demonstrations here, we imaged histone modifications of hundreds of genomic loci simultaneously. Since MERFISH allows >10,000 distinct RNAs to be imaged and identified in individual cells (Xia et al., 2019), we anticipate that the genomic throughput of epigenomic MERFISH could be further increased to allow simultaneous profiling of thousands of genomic loci with specific epigenetic modifications.

We further demonstrated that epigenomic MERFISH can be applied to tissue samples. Using this approach to spatially profile two distinct histone modifications that mark active promoters and putative enhancers, we observed region-specific distributions of active promoters and putative enhancers in both adult and developing mouse brain. These measurements not only showed spatial patterns of active promoters and enhancers that are consistent with known spatial patterns of gene expression and region-specificity of putative enhancers in the mouse brain, but also revealed previously unknown fine spatial distributions of putative enhancers as well as putative enhancer-promoter pairs and enhancer hubs for regulating genes involved in brain development.

Compared to sequencing-based single-cell epigenomic profiling methods that requires cell dissociation, epigenomic MERFISH retains the spatial context of cells and hence enables the spatially resolved single-cell profiling of epigenetic activities in tissues. In parallel to our work, a sequencing-based spatial epigenomic profiling method (hsrChST-seq) has been developed by performing CUT&Tag *in situ*, followed by microfluidics-assisted spatial barcoding and sequencing (Deng et al., 2021). Compared to the spatial resolution of hsrChST-seq (20 or 50 um pixel size), imaging-based epigenomic MERFISH has a much higher (sub-μm) spatial resolution, which not only facilitates single-cell analysis but should also allow sub-nuclear organization of the epigenome to be probed within individual cells. Epigenomic MERFISH also has its limitations. Unlike sequencing-based methods, which allow untargeted genome-wide detection of epigenetic sites, epigenomic MERFISH is a targeted approach and hence requires prior knowledge or hypothesis for the selection of epigenomic loci. We do, however, anticipate that it will be possible to profile the epigenetic properties of thousands, perhaps even tens of thousands, of genomic loci simultaneously, which should partially mitigate this limitation.

We foresee many possible applications of epigenomic MERFISH. Here, we demonstrated epigenomic MERFISH by spatially profiling histone modifications that mark active promoters and enhancers. The *in situ* tagmentation can be applied to capture other epigenetic marks, as long as antibodies or other affinity probes for these marks exist. Thus, we anticipate that epigenomic MERFISH can be applied to study many epigenomic properties, providing spatially resolved single-cell profiling of not only epigenetic (histone and DNA) modifications, but also the binding patterns of transcription factors, cofactors, and non-coding RNAs along the genomic DNA.

Recent studies demonstrated the possibility to preload PA-Tn5 with antibodies to target different epigenetic marks simultaneously (Gopalan et al., 2021). We envision that this approach can also be applied to epigenomic MERFISH, making it possible to simultaneous map multiple distinct epigenetic marks in the same cells, for example marking promoter and enhancer activities by measuring H3K4me3 and H3K27ac activities simultaneously to provide a more comprehensive picture of enhancer activity and gene regulation within the cell.

As an imaging approach, we also envision that epigenomic MERFISH can be combined with different modalities of imaging-based omics measurements, such as 3D genome and transcriptome imaging, to enable simultaneous detection of the epigenetic and protein-binding profiles of chromatin, the 3D organization of the chromatin, and the gene expression profiles within the same cells. Such a single-cell spatial multi-omics approach promises to substantially accelerate our understanding of the mechanisms underlying transcriptional regulation and the role of gene regulation in tissue development and functions.

## ACKNOWLEDGEMENTS

We thank Justyna Janas for providing initial aliquots of PA-Tn5. We thank Bogdan Bintu and Rongxin Fang, and other members of the Zhuang lab for helpful discussions. This work is supported in part by the National Institutes of Health (to X.Z.). T.L. is supported in part by the Harvard Biological and Biomedical Sciences Program. X.Z. is a Howard Hughes Medical Institute Investigator.

## AUTHOR CONTRIBUTIONS

T.L., A.C.E., and X.Z. designed the experiments. T.L., A.C.E. performed experiments and analyses. T.L., A.C.E, and X.Z. interpreted data and wrote the manuscript. X.Z. supervised the study.

## STAR Methods

### KEY RESOURCES TABLE

| REAGENT or RESOURCE | SOURCE | IDENTIFIER |
| --- | --- | --- |
| Antibodies |  |  |
| Anti-H3K27ac antibody | Abcam | Cat# ab4729, RRID: AB_2118291 |
| Anti-H3K4me3 antibody | Thermo | Cat# 49-1005, RRID: AB_2533856 |
| Donkey anti-Rabbit IgG (H+L) Highly Cross-Adsorbed Secondary Antibody, Alexa Fluor 488 | Thermo | Cat# A-21206, RRID: AB_2535792 |
| SON DNA Binding Protein (SON) (N-Term) antibody | antibodies-online | ABIN768615, RRID: AB_11207358 |
| Chemicals, Peptides, and Recombinant Proteins |  |  |
| Formamide | Ambion | Cat# AM9342 |
| 20xSSC | Ambion | Cat# AM9763 |
| Triton-X | Sigma | Cat# T8787 |
| Glucose oxidase | Sigma | Cat# G2133 |
| Phusion® Hot Start Flex 2X Master Mix | New England Biolabs | Cat# M0536 |
| HiScribe™ T7 Quick High Yield RNA Synthesis Kit | New England Biolabs | Cat# E2050 |
| Maxima H Minus Reverse Transcriptase | ThermoFisher | Cat# EP0752 |
| dNTP mix | ThermoFisher | Cat# R1121 |
| Eagle's Minimum Essential Media (EMEM) | ATCC | Cat# 30-2003 |
| Fetal Bovine Serum Plasma | Sigma | Cat# F1141 |
| 32% Paraformaldehyde | Fisher Scientific | Cat# 50-980-494 |
| MEGAscript Transcription Kit | Thermo | AMB13345 |
| aminoallyl-UTP | Thermo | R1091 |
| Yeast tRNA | Thermo | 15401029 |
| Stellaris RNA FISH hybridization buffer | Biosearch | SMF-HB1-10 |
| PA-Tn5 | Diagenode | C01070002 |
| 1M HEPES | Thermo | 15630080 |
| Spermidine | Sigma | S2626-5G |
| T4 DNA ligase | NEB | M0202L |
| T4 DNA polymerase | NEB | M0203L |
| DAPI | ThermoFisher | Cat# D1306 |
| RNase inhibitor, Murine | New England Biolabs | Cat# M0314 |
| RNasin™ Plus RNase Inhibitor | Promega | Cat# PRN2615 |
| 1M Tris, pH 8 | ThermoFisher | Cat# 15568025 |
| Catalase | Sigma | Cat# C3155 |
| 6-hydroxy-2,5,7,8-tetramethylchroman-2-carboxylic acid (Trolox) | Sigma | Cat# 238813 |
| tris(2-carboxyethyl)phosphine (TCEP) | Sigma | Cat# C4706 |
| UltraPure™ BSA (50 mg/mL) | Thermo | Cat# AM2618 |
| Experimental Models: Cell Lines |  |  |
| hTERT-RPE1 | ATCC | Cat# CRL-4000 |
| P56, C57BL6 male mice | Charles River | Cat# 027 |
| Time-pregnant mice E13.5 | Charles River | Cat# 027 |
| Oligonucleotides |  |  |
| Primers, readout probes | Integrated DNA Technologies | See Table S1 |
| Encoding oligonucleotide probe libraries | Twist Bioscience | See Tables S1 |

### RESOURCE AVAILABILITY

#### Materials Availability

Oligonucleotide probe sequences used for imaging can be found in Table S1. These probes or templates for making these probes can be purchased from commercial sources, as detailed in the Key Resources Table.

#### Data and Code Availability

Sequencing data have been deposited to NCBI GEO data repository (GSE191069). All data reported in this work are available upon request. Analysis software used in this work is available at https://github.com/TianLuHarvard/Code.

### EXPERIMENTAL MODEL AND SUBJECT DETAILS

#### Cell lines

hTERT-RPE1 cells (ATCC, CRL-4000) were cultured in DMEM/F-12, GlutaMAX™ supplement (Thermo, 10565042), 10% FBS (Sigma, F4135-1L) and 1% Pen/Strep (Invitrogen, 15140122) antibiotics at 37 °C. ∼1 million cells were plated onto silanized coverslips one day before fixation.

#### Mouse brain tissue sections

P56 male C57BL6 mice were ordered from Charles River Laboratories. The brain was dissected after euthanasia and embedded in OCT (VWR, 25608-930) and stored at −80°C. E13.5 embryonic brains were collected from timed pregnant C57BL6 mice ordered from Charles River Laboratories. The brain was dissected from the embryo and embedded in OCT and stored in −80°C. The embedded tissue was transferred to −18°C cryostat before slicing. The tissue was sliced into 10μm thick tissue slices and mounted onto silanized coverslips. The mounted tissue was left at room temperature for at least 10 mins before fixation. Animal care and experiments were carried out in accordance with NIH guidelines and were approved by the Harvard University Institutional Animal Care and Use Committee (IACUC).

### METHOD DETAILS

#### Oligonucleotide Probe Design

##### Selection of target epigenetic loci

The candidate target epigenetic loci for each of the experiments were selected using the criteria as described below. Those candidate loci were subsequently filtered using criteria listed in the *Encoding probe design* section.

1. H3K27ac loci in hTERT-RPE1 cells H3K27ac peaks were obtained by performing the CUT&Tag reaction in house following the published protocol (Kaya-Okur et al., 2019). The sequencing reads were mapped with *bowtie2*, followed by peak calling using *MACS* with q<0.10 and consistent peaks between two replicates were identified by performing irreproducible discovery rate (IDR) setting the threshold at q<0.10 (see Sequencing Data Analysis for more information).
2. H3K4me3 loci for essential genes in hTERT-RPE1 cells H3K4me3 marks at the promoter regions (±2kb of TSS) were used to estimate the detection efficiency of epigenomic MERFISH. The promoters of housekeeping genes (*Gapdh, Bactin*) and highly expressed and essential genes (Hart et al., 2015) were selected.
3. H3K4me3 loci in adult mouse cortex and embryonic mouse brain Candidate loci that potentially show layer-specific enrichment pattern in adult mouse cortex were curated using marker genes reported by Loo and coworkers (Loo et al., 2019). The gene list included marker genes for layer II-VI excitatory neurons, inhibitory neurons, microglia, endothelial cells, astrocytes, and oligodendrocytes. Candidate loci that show region specific enrichment pattern were chosen as a list of homeodomain transcription factors with well characterized expression in the embryonic brain.
4. ATAC-seq peaks in adult mouse cortex Candidate ATAC-Seq peaks were curated from the data published (Gray et al., 2017). There were four mCherry sorted populations of cells driven by the following gene promoters: Cux2, Ntsr1, Rbp4 and Snnc1 which represent layers II/III, IV, V and VI excitatory neurons respectively. Also, we selected peaks that are more than 2 kb away from TSS to avoid mapping the promoter associated H3K27ac peaks. We also required that the selected ATAC-Seq peaks has >0 H3K27ac reads mapped to it, according to the bulk H3K27ac ChIP-Seq data obtained from sorted CAMKII excitatory neurons (Mo et al, 2015). Only peaks that differentially expressed (differential binding performed in diffBind, FDR p value<0.05) in each cortical layer were included in the peak selection.
5. H3K27ac ChIP-seq peaks in the embryonic mouse brain E15.5 embryonic fore, mid, and hindbrain specific H3K27ac ChIP-seq peaks previously identified by the ENCODE consortium were merged using *bedTools merge* to obtain a master list of fore, mid, and hindbrain peaks (Gorkin et al., 2020). The read counts for the master list of peaks were calculated using *featureCount* with default settings. The read counts were subsequently normalized using *DESeq2 norm*. Adjusted p-values were obtained using DESeq2 by doing pairwise comparison between any two brain regions. Peaks that were significant (adj p-value <0.05) against the other two brain regions and more than 4-fold differentially expressed and with normalized reads number >200 were included as candidate loci. 5 loci that have less than 20 reads in H3K27ac ChIP-Seq data in fore, mid, and hindbrain were additional included as control loci. These control loci served a negative control for our epigenomic MERFISH experiment.

##### Encoding probe design

During epigenomic MERFISH imaging, a library of encoding probes was first added to the sample to bind to the RNAs generated by *in situ* T7 transcription. These encoding probes each has a 30- nt target sequence that can bind to a 30-nt target region on one of the RNAs, and 3 readout sequences that allows the encoding probes to be detected by complementary fluorescently labeled readout probes. Each distinct readout sequence corresponds to one bit in the barcode and the collection of readout sequences on an RNA determines the barcode of the RNA. For example, if the barcode reads “1” at bits 1, 3, 5, and 7 and “0” at all other bits, the collection of encoding probes on the RNA should contain readout sequences 1, 3, 5 and 7.

To design encoding probes that target human or mouse loci respectively, hg19 and mm10 genome builds were used for designing target sequences on the encoding probes. For each locus, we identify a number of 30-nt long target regions for the MERFISH encoding probes to bind. The candidate target regions were selected from a sliding window of 30 nt starting from the first nucleotide. The candidate target regions were kept if

1. The range of GC% 33%-73% and Tm 61°C-81°C
2. It doesn’t have same 15 nt sequences within other ChIP-seq and IgG peaks
3. It doesn’t contain more than 3 consecutive dinucleotide repeats

The next candidate target region was chosen such that it can have a <4 nt overlap with the previous target region and was on the other strand of the DNA. This was repeated until the whole candidate locus was covered. This strategy allows RNA transcribed from either ends of the tagged DNA fragment to be imaged by MERFISH. After designing probes for all candidate loci, the list of candidate loci in the library were filtered by the requirement that number of encoding probes for each locus was > 1.2-2 probes per 100 bp in order to ensure efficient labelling of different RNA lengths.

Template library for synthesizing encoding probes were purchased from Twist Bioscience. The sequences for encoding probes are listed in Table S1.

##### Readout probes

Dye-labeled readout probes were purchase from Integrated DNA Technologies. In the readout probes, fluorescent dye molecules (Alexa 750, Cy5, or Atto565/Cy3B) were linked to oligonucleotide via a disulfide bond that can be cleaved by TCEP. The sequences for dye labeled readout probes are listed in Table S1.

##### Barcode design

The 24-bit Hamming distance 4 (HD4) and Hamming weight 4 (HW4) code, which contains 366 distinct barcodes, were adopted from La Jolla covering repository and used for MERFISH imaging in this work. The barcodes were then randomly assigned to the target loci except for the requirement that for each bit, there were only 3-5 “on” bits (bits that read ‘1’) for each chromosome, in order to ensure that the number of spots imaged in each bit were sufficiently sparse.

After barcode arrangement, we next assigned the readout sequences to the encoding probe. Since the HW4 code contains four “1” bit per barcode, 4 readout sequences were assigned to each locus. We required that the three readout sequences on each encoding probe correspond to three of the four readout sequences assigned to its target locus and any two adjacent encoding probes have all four readout sequences. This strategy aims to let short RNA that can only fit 2 probes have all 4 bits presented on the encoding probes.

#### Experimental Setup

##### Microscope setup for image acquisition

We used two microscope setups to perform the imaging, and the setups were as described previously (Moffitt et al., 2016a; Wang et al., 2019a). In one of the setups, a Nikon CFI Plan Apo Lambda 60× oil NA 1.4 immersion objective installed on a Nikon Ti-U microscope body was used for imaging. Illumination was provided by solid-state single-mode lasers (405 nm laser, Obis 405 nm LX 200 mW, Coherent; 488 nm laser, Genesis MX488-1000, Coherent; 560 nm laser, 2RU-VFL-P-2000-560-B1R, MPB Communications; 647 nm laser, 2RU-VFL-P-1500-647-B1R, MPB Communication; and 750 nm laser, 2RU-VFL-P-500-750-B1R, MPB Communications). Mechanical shutters were used to switch the 750 nm laser. Acousto-optic tunable filters (AOTF) were used to control the intensities of the 488 nm, 560 nm, and 647 nm lasers; the 405 nm laser was modulated by a direct digital signal. To separate the excitation illumination from the fluorescence emission, a custom dichroic (Chroma, zy405/488/561/647/752RP-UF1) and emission filter (Chroma, ZET405/488/461/647-656/752m) were used. The emission was imaged onto the Hamamatsu digital CMOS camera. During acquisition, the sample was translated using a motorized XY stage (Ludl, BioPrecision2) and kept in focus using a home-built autofocus system. A peristaltic pump (Gilson, MINIPULS 3) pulled liquid into Bioptechs FCS2 flow chamber with sample coverslips and three valves (Hamilton, MVP and HVXM 8-5) were used to select the input fluid.

In the other setup, samples were imaged on a custom-built Olympus microscope body. Laser illumination was provided at 750, 647, 560, 488 and 405 nm with a Lumencor Celesta light system. These illumination laser wavelengths were used to excite Alexa750, Cy5 and Cy3 conjugated readout probes, Alexa-488 fiducial beads and DAPI respectively. The setup for the rest of the imaging system was as described previously.

#### Experimental Procedures and Protocols

##### Encoding probe synthesis

Encoding probe synthesis was as described previously (Xia et al., 2019). Briefly, the library was ordered from Twist Bioscience and diluted in TE buffer to about 1 ng/µL. We did qPCR (10-12 cycles) to amplify the oligo pools and stopped the reaction when the curve started to plateau. The amplified templates were purified and transcribed into RNAs via *in situ* transcription for >20 hrs at 37°C, The RNAs were reverse transcribed to ssDNAs for 1hr at 55°C, and we then purified the DNAs via alkaline hydrolysis (to remove RNA templates), phenol-chloroform extraction (to remove proteins), and ethanol precipitation (to remove nucleotides and concentrate probes). The final concentration of the encoding probe library was about 40,000 ng/µL,

##### Imaging coverslip silanization

40-mm, round #1.5 coverslips (Bioptechs, 0420-0323-2) were first cleaned by 37.5% HCl and pure methanol for 30 mins at room temperature, washed by 70% ethanol and dried. For silanization, coverslips were covered in silanization buffer (500 mL distilled water, 1500 µL Bind-silane (Sigma, GE17-1330-01) and pH adjusted to 3.5 by glacial acetic acid) for an hour at room temperature. The coverslips were then washed with water and dried in the oven before storing in a dehumidified chamber.

##### Epigenomic MERFISH protocol in cell culture and tissue slices

For cell culture, the cells were fixed with 1% PFA in 1x PBS for 5 mins at room temperature and washed three times with 1× PBS. The sample was then permeabilized by 1% Triton-X for 20 mins at room temperature and washed three times with 1× PBS. Then 0.1 M HCl was added for chromatin loosening for 5 mins at room temperature and washed three times with 1× PBS. The sample was then incubated in block buffer (50% Ultrapure BSA, 1% Triton-X and 1× PBS) for one hour at room temperature and further incubated in 1:100 primary antibody (against H3K27ac or H3K4me3) for one hour at room temperature. The sample was washed three times with 1x PBS and incubated in 1:200 secondary antibody (Thermo, A-21206) for one hour at room temperature. After washing three times with 1x PBS, the sample was ready for transposition.

Before the transposition, PA-Tn5 (Diagenode, C01070002) was loaded with a pair of annealed loader DNAs ordered from IDT (sequences in Table S1). The two loader DNAs were annealed at 100 μM concentration separated using the following settings at the thermocycler (95°C for 6mins and −5 °C per cycle for 15 cycles). For loading PA-Tn5, 6.5 μL of annealed loader A, 6.5 μL of annealed loader B and 10 μL of PA-Tn5 were mixed and incubated at room temperature for one hour. 12.5 μL of 100% glycerol was added to the mixture and PA-Tn5 was ready for transposition and stored at −20°C.

The 50 mL PA-Tn5 binding buffer contained 1 mL 1M HEPES (Thermo, 15630080), 3 mL 5 M NaCl, 4 μL Spermidine (Sigma S2626-5G), 100 µl of 5% digitonin, and water filled up to 50 mL. The sample was incubated with 1:50 PA-Tn5 in 50 μL of PA-Tn5 binding buffer at room temperature for 1 hour to let protein A bind to antibodies. The high salt concentration in the buffer prevents the non-specific binding of the PA-Tn5 and the lack of Mg^2+^ in the buffer prevents the transposition of PA-Tn5 (Kaya-Okur et al., 2019). After the incubation, samples were washed with PA-Tn5 binding buffer 3 times to remove nonspecific binding. Samples were then incubated in the PA-Tn5 transposition buffer (1 mL of PA-Tn5 binding buffer with 10 μL 1M MgCl_2_) for 1hr at 37°C for transposition.

After transposition, samples are washed three times with 1× PBS and then embedded in 4% Acrylamide/Bis 19:1 gel with 1:200 Alexa-488 beads (Invitrogen) for 1hr at room temperature. The embedded sample was digested in 2% SDS, 0.5% Triton-X and 1:100 proteinase K in 2x SSC at 37 °C for at least 16 hours. The sample was then washed 3 times with 1× PBS. Each wash was one hour at room temperature on a shaker. After wash, the sample was incubated in 50 μL nick ligation mix containing 2.5 μL of T4 DNA ligase (NEB, M0202L), 2.5 μL of T4 DNA polymerase (NEB, M0203L), 5 μL of T4 DNA ligase buffer (NEB, M0202L), 5 μL of 10 mM dNTP (NEB, N0447L) and 35 μL of water at room temperature for 40 minutes at room temperature. After wash three times with 1× PBS, the sample was incubated in transcription mix (MEGAscript Transcription Kit; ThermoFisher, AMB13345) with aminoallyl-UTP (Thermo, R1091) at 37°C for 16-18 hours. The 200 μL transcription mix contained 20 μL ATP, 15 μL UTP, 20 μL CTP, 20 μL GTP, 20 μL 10x Reaction buffer, 20 μL T7 polymerase, 10 μL Rnase inhibitor and 5 μL aminoallyl-UTP. Samples for RNA sequencing didn’t have aminoallyl-UTP. After *in situ* transcription, the samples were fixed in 4% PFA for 20 mins at room temperature to crosslink the transcribed RNA via the aminoallyl group to the gel and stained with 1:30 encoding probe library in 30% hybridization buffer at 37°C overnight. The hybridization buffer contained 30% formamide (ThermoFisher, AM9342), 60% stellaris RNA FISH hybridization buffer (Biosearch, SMF-HB1-10), 10% 25 mg/mL Yeast tRNA (ThermoFisher, 15401029), and 1:100 murine RNase inhibitor.

For tissue slides, the protocol was similar, except that 1) the primary antibody was stained overnight at 4°C, 2) the binding of the PA-Tn5 took 2 hrs and the tagmentation was incubated overnight at 37°C.

For SON test in Figure S1, the samples were stained with SON antibodies (antibodies-online, ABIN768615) and the PA-Tn5 was loaded with a pair of Tn5 loader DNAs (MEA and MEB) from Illumina, and the loader DNAs contained a single-stranded overhang. After tagmentation, the samples are washed with PBS and hybridized with 100 nM probes with a region that can target the overhang region and another that can bind to readout probes in 10% hyb buffer (10% formamide, 80% stellaris RNA FISH hybridization buffer, 10% 25 mg/mL Yeast tRNA, and 1:100 murine RNase inhibitor) at 37°C for 1 hr. The samples were then washed in 30% formamide with 2x SSC and stained with readout probes in 10% EC buffer for imaging.

##### RNA extraction and sequencing

The RNAs were harvested from the polyacrylamide gel using the crush and soak method. Briefly, the gel was scraped off the coverslip surface and shredded into tiny pieces before resuspension in the elution buffer (500mM ammonium acetate and 1mM EDTA-KOH pH 8.0) with 1:100 RNase inhibitor (Promega) and rotated at room temperature for 2-4 hrs. After 2-4 hours, gel pieces were removed using a 40um filter. RNAs in the eluate were then purified using Zymo RNA kit using the recommended protocol by Zymo. RNAs were first reverse transcribed into cDNAs using a primer specific to one mosaic end using Maxima RT kit (ThermoScientific) using the manufacturer’s protocol. The resulting cDNAs were PCR enriched using the following settings using Phusion (NEB). The number of cycles (N) were first determined by running a test PCR reaction (¼ the saturation). The usual cycle number required was around 12-15.

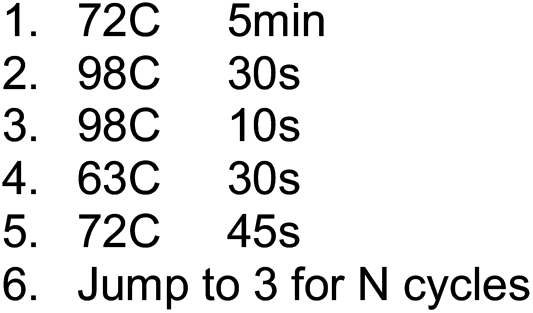

Following PCR, 0.7 volume of AMPure XP beads (Beckman) was added to the PCR reaction. The mixture was incubated for 5 minutes before placing on the magnetic rack. The beads were then washed twice with 80% ethanol before they were let dry at room temperature. 20ul of RNase free water was added for elution. The 0.7 volume ensured proper removal of the primer dimer peak. Subsequently, the PCR product was examined using DNA Tapestation for proper size distribution. Concentration of the PCR product was obtained by selecting the peak from 100-1000bp. Libraries that passed the Tapestation QC were sequenced paired-ended using Novaseq SP100 or Nextseq platform.

##### Imaging procedure for epigenomic MERFISH

After hybridization of the encoding probes, samples are washed in 30% formamide at room temperature for 20 minutes and washed 3 times with 2x SSC before imaging. The first set of readout probes were added at 3 nM concentration in 2X SSC with 10% ethylene carbonate. The stained sample coverslips were mounted to the Bioptechs imaging chamber for imaging. Each imaging round contained three distinct steps: imaging, cleave and hybridization. The buffer for each step was flowed into the imaging chamber via a fluidic system controlled by a custom made software (Su et al., 2020).

In the imaging step, about 2 ml of anti-photobleaching buffer was flowed into the chamber. For anti-photobleaching buffer, we used either

1. rPCO-PCA based buffer: 2x SSC, 5 mM 3,4-dihydroxybenzoic acid (Sigma, P5630), 2 mM trolox (Sigma, 238813), 50 µM trolox quinone, 1:500 rPCO (Oriental Yeast Company), 1:500 Murine RNase inhibitor, and 5 mM NaOH (to adjust pH to 7.0) and topped up to 50ml with nuclease free water.
2. Glucose oxidase base buffer: 50 mg glucose-oxidase (Sigma, G2133), 50 mg (±)-6-Hydroxy-2,5,7,8-tetramethylchromane-2-carboxylic acid (Trolox) (Sigma, 238813), 300 µL catalase (Sigma, C100-500MG), 10% w/v glucose (Sigma, G8270), 5 mL 500 µM Trolox quinone and 50 µL murine RNase inhibitor) and topped up to 50ml with nuclease free water.

For cultured hTERT-RPE1 cells, 100 fields of view were imaged for each sample with 40 z-planes (step size of 200 nm) imaged per channel. For brain tissue slices, 200-400 fields of view were imaged for each sample with 30 z-planes (step size of 300 nm) per channel. The images were acquired at 10 Hz. After imaging, in the cleaving step, 2 ml of cleaving buffer containing 2x SSC and 50 mM TCEP (Goldbio) was flowed into the imaging chamber to cleave the dye off the readout probes and left for 12 minutes before the residual TCEP was removed by flowing in 2ml of 2x SSC.

In the readout hybridization step, readout probes in three different colors (labeled with Alexa750, Cy5, and Atto565/Cy3B respectively) were added to the hybridization buffer (2x SSC, 10% ethylene carbonate, 200μl of 100% Triton-X) at a concentration of 3 nM for each readout probe. The readout probes were left to hybridize for 12 minutes before the unbound readout probes were washed away with a wash buffer (2x SSC, 10% ethylene carbonate). The three steps were repeated 8 times for a 24 bit imaging.

#### Image Analysis

##### Decoding of Epigenomic MERFISH spots

To normalize for intensity variation across different color channels, every image in a given color channel was divided by the mean-intensity image of all images in that that color channel. Images of multiple rounds were registered using Alexa 488 fiducial beads. Cell nuclei were segmented by watershed algorithm using DAPI staining as both seed and boundaries.

Epigenomic MERFISH signals from each channel in each hybridization round were identified using two spot-finding methods: In the experiments in which we segmented individual cells, the pixels in each nucleus with intensity higher than certain brightness threshold was selected. In order to connect spots detected in different z-planes, the selected pixels across different z planes were clustered by the *bwareaopen* function in MATLAB with the requirement that the number of pixels within a cluster should have in the range of 10-100. In order to capture clusters with relatively wide variations in spot intensity, this process was iterated using multiple brightness thresholds (from top 0.001% to top 1% of the FOV with the decrement of 0.01%). Each iteration of lowering the brightness threshold allowed the identification of additional clusters that belonged to one of the following two types: (I) dim pixel clusters that could not be recognized at higher brightness threshold in the previous iteration, and (II) larger clusters that encompassed one or more pixel clusters found in the previous iteration. Any cluster of type-I was preserved only if its total number of pixels is within the range of 10-100. If the pixel number of any cluster of type-II fell within the permissible range, it was kept; if not, it was deleted, and the smaller pixel cluster(s) found in the previous round that overlapped with this new cluster were kept instead. Each pixel cluster was then considered a spot and the x, y, and z coordinates of the spot was measured using the *regionprops3* function in MATLAB.

In the experiments in which we didn’t segment the cells, the 3D spot finding approach described above was computationally too slow to find the spots in the whole imaging field of view. We thus used a 2D spot finding approach first. Briefly, the 2D spots for each Z plane were first identified using the approach described above, but in 2D instead of 3D, using the *regionprops* function. After spot finding in every z plane, the 2D spots across all z planes were clustered by *DBSCAN* using the distance threshold of 50 nm in the x, y plane and 300 nm in z plane and minimum spot number of 2. Each resulting spot cluster was then considered a 3D spot and the x, y, and z coordinates of the 3D spot were calculated as the mean of x, y, and z coordinates of the 2D spots in the spot clusters.

The spots identified from two methods were further filtered by signal-to-background ratio with a threshold of 1.4. The signal-to-background ratio for a spot was defined as the intensity of the center of the spot divided by the minimal intensity of the pixels that were 500 nm away from the spot center in the xy plane.

After 3D spot finding, the spots from all bits were clustered by *DBSCAN* using a threshold of minimal distance of 250 nm in 3D and minimum spot number of 3. For any cluster that had 3-5 spots (i.e. a cluster that was detected in 3-5 bits), the cluster was then decoded according to the codebook allowing at most one-bit mismatch from the valid barcodes. For any cluster that had more than 5 spots, the spots within that cluster were further clustered by DBSCAN using a threshold of minimal distance of 150 nm in 3D and minimum spot number of 3 and the resulting new clusters that had 3-5 spots were decoded. The clusters that were not matching to any barcode were discarded. The final x, y, and z coordinates of the decoded spots were calculated as the mean x, y, and z coordinates of the spot across all bits. The decoded locus identity, 3D localization and barcode error of the spots are saved for further analysis.

### QUANTIFICATION AND STATISTICAL ANALYSIS

#### Quantifying region-specific enrichment in mouse embryonic brain

The specific regions of the brain were manually segmented by comparing the DAPI staining in our images and the reference Allen brain atlas. The total number of decoded spots within these regions were counted and divided by the DAPI positive area to calculate the spot density. The ordering of the heatmaps for imaging data in Figures 3 and 5 were done as follows: 1) The loci with maximum density in certain regions were grouped. 2) Within this group, the maximum density for those loci were ordered from largest to smallest. 3) Arrange the group in the region order as shown in the figures. All heatmaps were plotted using the Z-score of the spot density for each locus.

#### Quantifying layer-specific enrichment in adult mouse cortex

The layers in the mouse cortex were manually identified by comparing the DAPI staining in our images and the reference Allen brain atlas. To assign each cell into each layer, we approximated the cortical layer boundaries as a set of concentric circular arcs, which matched with the manually segmented cortical layer boundaries by visual inspection. We then determined whether any given cell belongs to a cortical layer by comparing the radial position of the cell with the radii of the concentric circular arcs representing the cortical layer boundaries. The layer enrichment for a specific epigenetic activity of each locus was calculated as follows: A cell was considered H3K4me3-positive (or H3K27ac-positive) for a certain locus if at least one decoded spot for this locus was detected in the cell. For each locus, the standardized layer enrichment in a specific layer was calculated as the z-score of the following quantity: the fraction of cells in the layer that were H3K4me3-positive (or H3K27ac-positive) for this locus. The significance of the enrichment was calculated using a chi-square test using *Chi2test* function in MATLAB. The ordering of the heatmaps for layer enrichment in Figures 2 and 4 were done as follows: 1) The loci with maximum layer enrichment in certain layers were grouped. 2) Within each group, the maximum layer enrichment for those loci were ordered from largest to smallest. 3) Arrange the groups in the layer order as shown in the figures. All heatmaps were plotted using the Z-score of the layer enrichment for each locus.

#### Clustering of putative active enhancers based on spatial distribution

The putative active enhancer loci marked by H3K27ac in the embryonic mouse brain were clustered based on their spatial distributions, measured as the number of decoded spots in each field of view (FOV). *Clustergram* function in MATLAB was used and the linkage for clustering the FOV and enhancer loci was ‘weighted’, distance was ‘Euclidean’. To identify the main clusters with more than 3 loci, we used the linkage threshold of 22.3. The resulting six main clusters were shown in the Figure 5.

#### Motif enrichment analysis

Motif enrichment analysis was performed by obtaining the sequences of the loci belong to each of the six main clusters described above using *bedtools getFasta*. The sequences were uploaded to the MEME-ChIP website and motif enrichment was performed for each of the six clusters using default settings. Only the top two most significant motifs were listed.

#### Sequencing data analysis

Sequencing reads were aligned to the hg19 and mm10 reference sequences using Bowtie 2.1.041. Mitochondrial reads were then removed. Reads were subsequently deduplicated using the rmdup option in samtools. Peak calling was performed using MACS 2.1.142 using the default options outlined in the vignette with the significant value cut-off at q<0.1. For experiments with two replicates, top 100,000 reproducible peaks were sorted by the p value and selected using IDR 2.0.243 with cutoff of IDR ≤ 0.1 using the recommended settings. Peaks were considered differentially expressed if they have q<0.10 by DiffBind. The global profiles of histone marks were then plotted using deeptools suite following the vignette centering at the peak summit.

## Supplementary Figures

**Figure S1.**
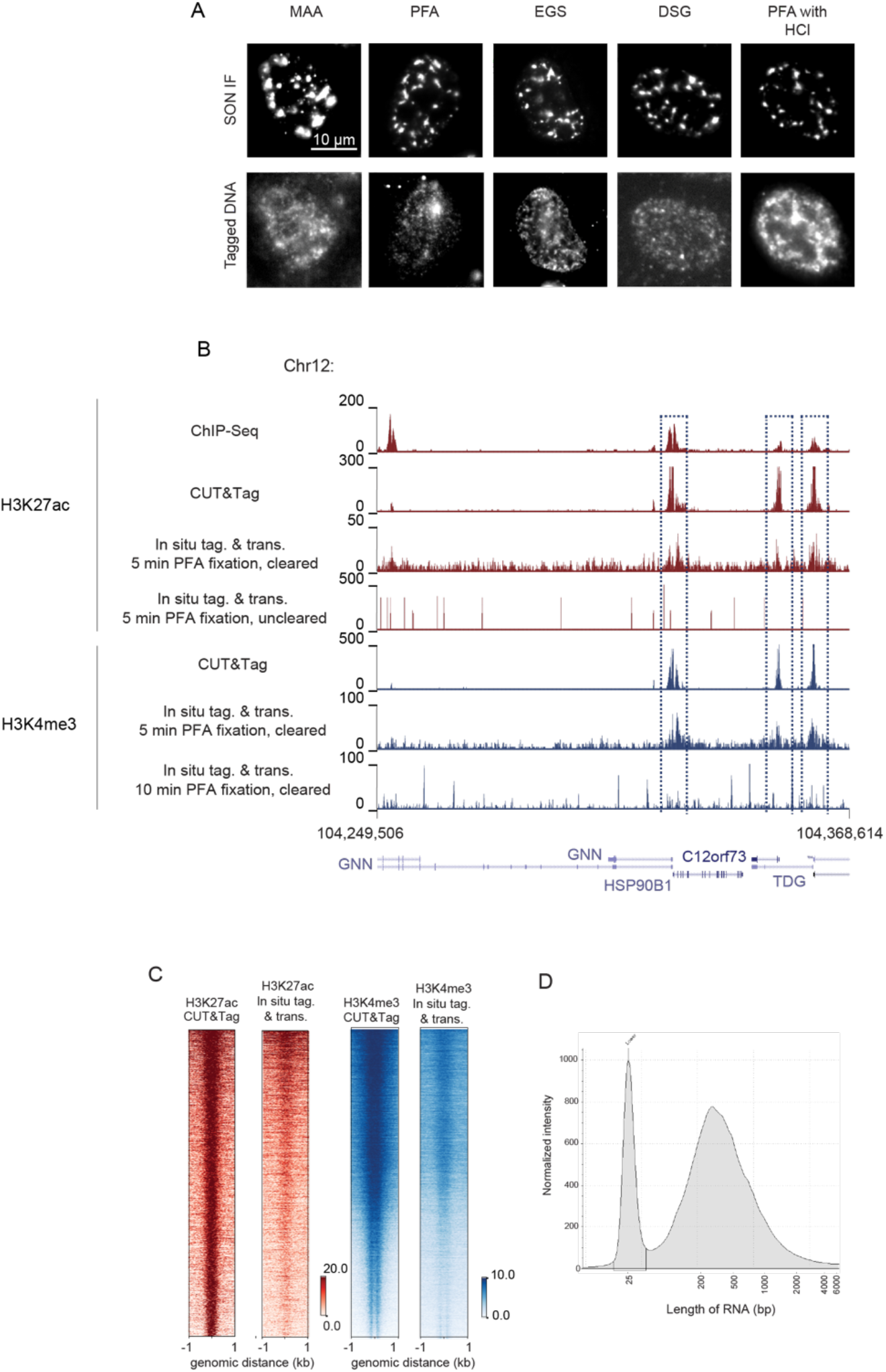
The effect of fixation and clearing on epigenomic MERFISH signals. (A) Images comparing the SON (a marker for nuclear speckles) immunofluorescence images (top row) in different fixation conditions [MAA=Methanol/acetic acid, 1%PFA, EGS=ethylene glycol bis(succinimidyl succinate), DSG=disuccinimidyl glutarate, 1%PFA followed by 0.1N HCl denaturation] and the location of tagged DNA fragments colocalized with SON and detected by FISH (bottom row). To generate tagged DNA fragments colocalized with SON, primary antibody against SON was added, followed by secondary antibody and PA-Tn5. 1%PFA with 0.1N HCl treatment gave the best colocalization between the SON immunofluorescence and the tagged DNA signals. We performed screening for fixation conditions using the nuclear speckle marker as it forms distinct puncta in the nucleus, enabling rapid screening of the *in situ* tagmentation conditions for fixed cells. (B) The first four of the UCSC tracks compare the sequencing results of *in situ* transcribed RNAs from tagged DNA fragments with the H3K27ac modification (*in situ* tag. and trans.) for cleared (5 min PFA fixation, cleared) and uncleared (5 min PFA fixation, uncleared) samples with results obtained from H3K27ac ChIP-Seq and CUT&Tag measurement in hTERT-RPE1 cells. The sequencing result of *in situ* transcribed RNA from cleared sample shows more unique reads and better capture of the epigenetic profile. The fifth, sixth and seventh UCSC tracks compare the sequencing result of *in situ* transcribed RNAs from tagged DNA fragments with the H3K4me3 modification (*in situ* tag. and trans.) obtained under two fixation conditions (5 min PFA fixation and 10 min PFA fixation) with H3K4me3 CUT&Tag in hTERT-RPE1 cells. The sequencing result of *in situ* transcribed RNA from 5 min fixation showed more unique reads and better capture of the epigenetic profile. The tracks were normalized to 1x sequencing depth. (C) Genome wide binding profiles comparing the sequencing results of *in situ* transcribed RNAs from tagged DNA fragments and the CUT&Tag results for the two chromatin marks (H3K4me3 (blue) and H3K27ac (red)) centered at the peak summits. Each pixel line represents a peak in CUT&Tag track and the same peak obtained from the sequencing result of *in situ* transcribed RNAs. The genomic locations of peaks are determined from the CUT&Tag data and the tracks for the *in situ* transcribed RNAs from the same genome regions are plotted for comparison. The scale bar shows the magnitude of the normalized peak height (1x sequencing depth). The peaks shown are significant peaks with q value < 0.1. (D) RNA Tapestation showing the RNA length distribution of in *situ* transcribed RNAs from tagged DNA fragments with the H3K4me3 modification.

**Figure S2.**
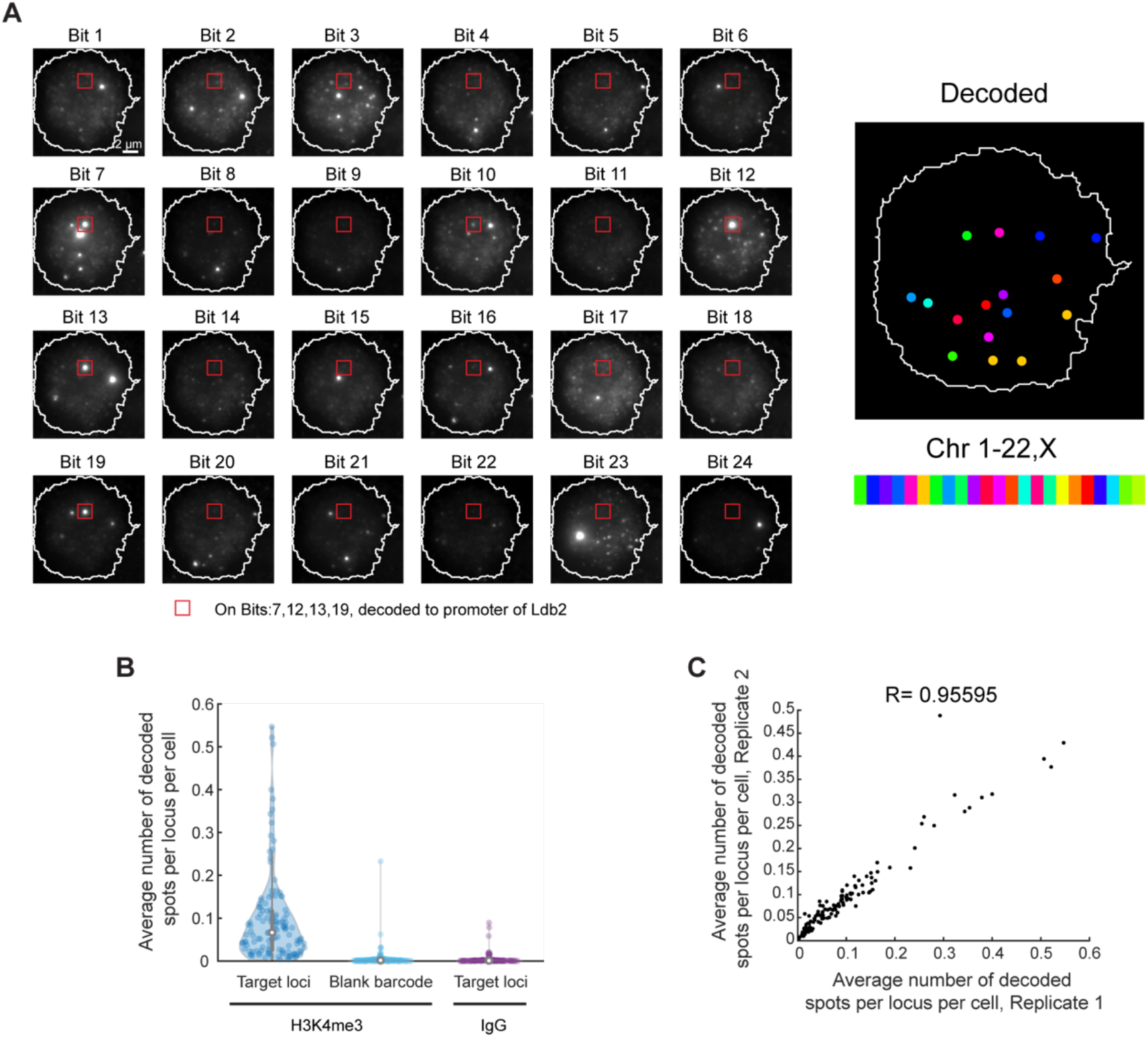
Accuracy, specificity, and reproducibility of epigenomic MERFISH measurements on mouse brain tissues. (A) Epigenomic MERFISH image of 127 target H3K4me3 loci in a single cell. The images from individual bits are shown on the left. The decoded image is shown on the right, with individual spots color-coded based on the identities of the chromosome identity. The spot, marked by the red box, was observed in bit 7,12,13,19 and hence decoded to the Chr5:44,797,746-44,801,746 locus, the promoter of *Ldb2*. (B) Violin plot showing the average number of decoded spots per cell for each target H3K4me3 locus (left) and each blank barcode (middle) when H3K4me3 antibody is used to capture the epigenetic mark. Also shown is the violin plot of the average number of decoded spots per cell for each target H3K4me3 locus when a control IgG is used instead (right). Each dot in the violin plots correspond to a single H3K4me3 locus or a blank barcode. (C) Scatter plot showing the correlation between two biological replicates of H3K4me3 imaging. Each dot corresponds to a single H3K4me3 locus.

**Figure S3.**
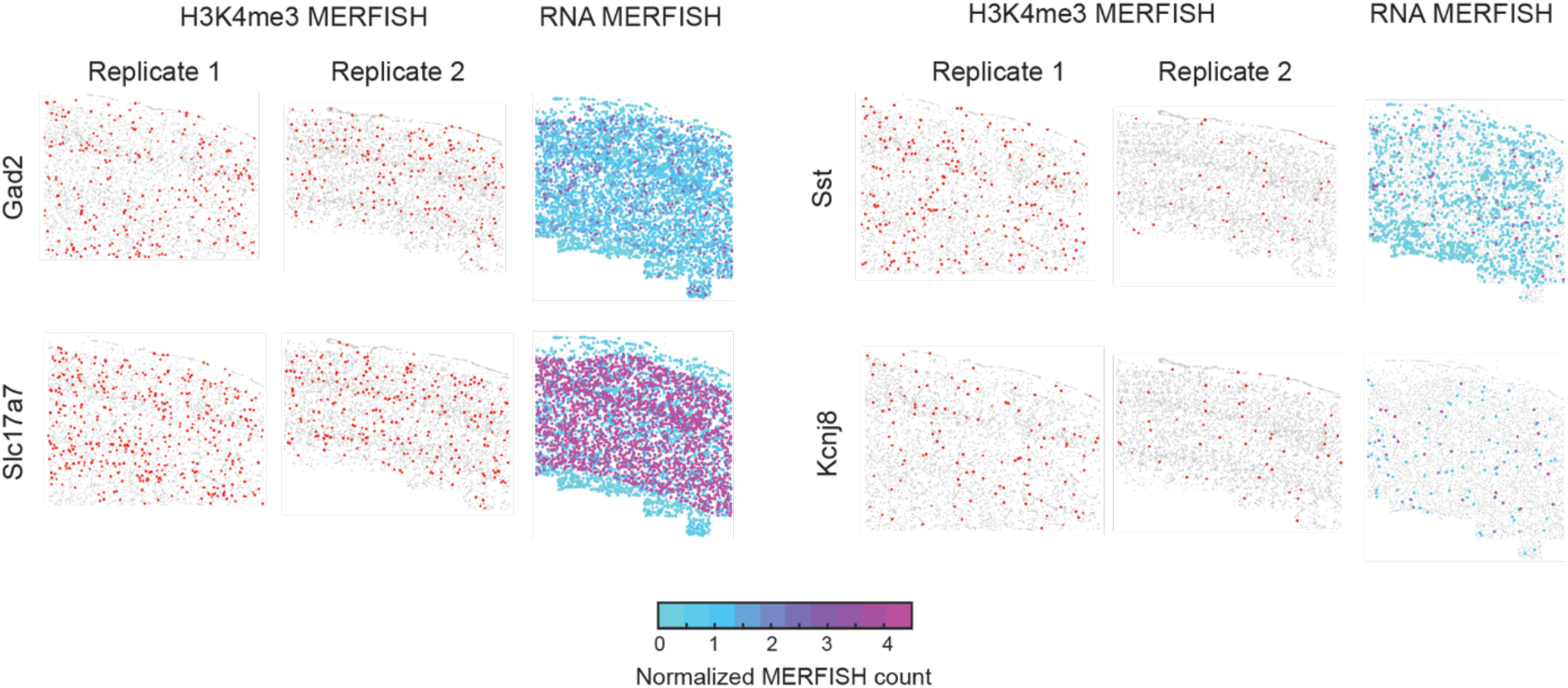
Epigenomic MERFISH images of several active promoters in mouse cortex and the RNA MERFISH images of corresponding genes. For each gene, the left panels show epigenomic MERFISH images of H3K4me3 signals for two replicates with each dot in the images representing a cell and red dots represent cells with positive H3K4me3 signals, and the right panel shows the RNA MERFISH image of the corresponding gene (Zhang et al., 2021). The RNA expression level in each cell is color coded according to the colored scale bar shown at the bottom.

**Figure S4.**
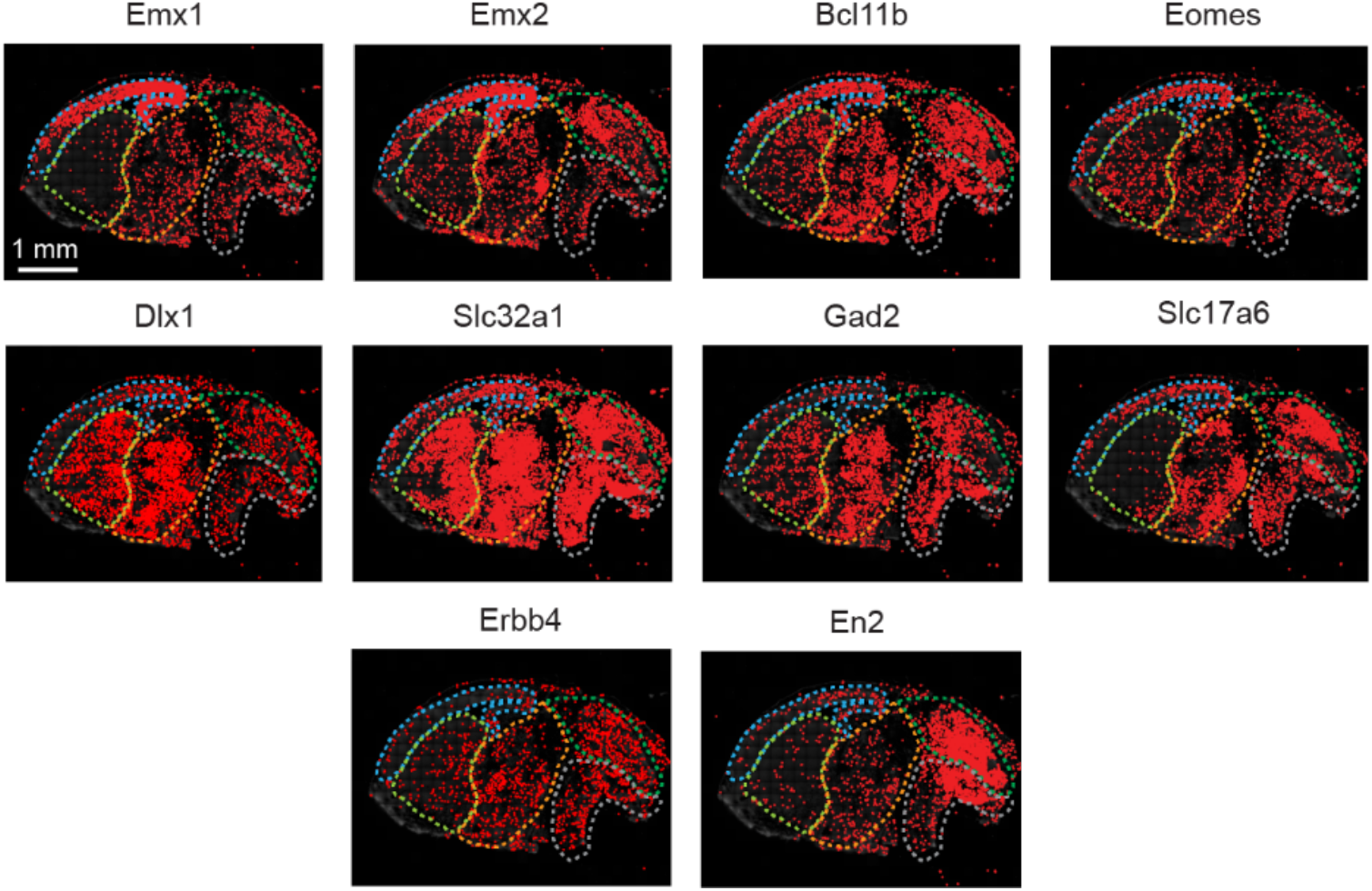
Epigenomic MERFISH images of additional representative active promoters in mouse embryonic brain. Epigenomic MERFISH images showing the H3K4me3 signals for ten additional promoters. The boundaries of cortex, subpallium, diencephalon, midbrain and hindbrain are marked by dashed line, according to the color scheme shown in Figure 3A.

**Figure S5.**
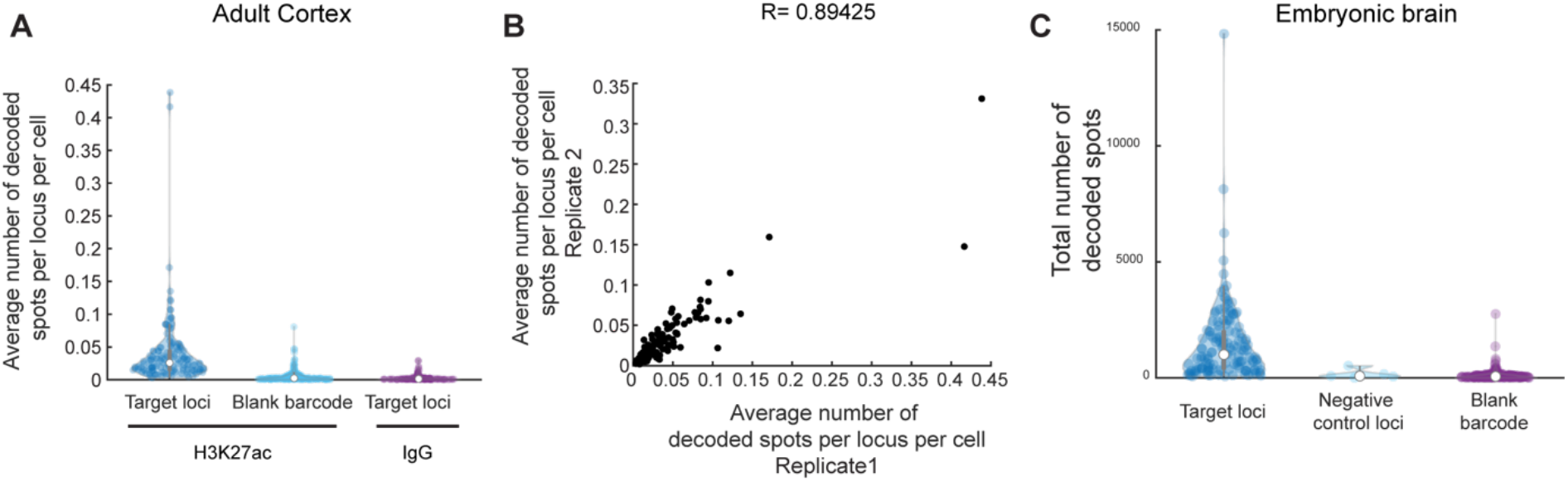
Accuracy, specificity, and reproducibility of epigenomic MERFISH measurements of target H3K27ac loci in mouse brain tissues. (A) Violin plot showing the average number of decoded spots per cell in adult mouse cortex for each target H3K27ac locus (left) and each blank barcode (middle) when H3K27ac antibody is used to capture the epigenetic mark. Also shown is the violin plot of the average number of decoded spots per cell for each target H3K27ac locus when a control IgG is used instead (right). Each dot in the violin plots correspond to a single H3K27ac locus or a blank barcode. (B) Scatter plot showing the correlation between two biological replicates of H3K27ac imaging in the adult mouse cortex. Each dot corresponds to a single H3K27ac locus. (C) Violin plot showing the total number of decoded spots in the embryonic mouse brain for each target H3K27ac locus (left), each negative control loci (middle) and each blank barcode (right).

**Figure S6.**
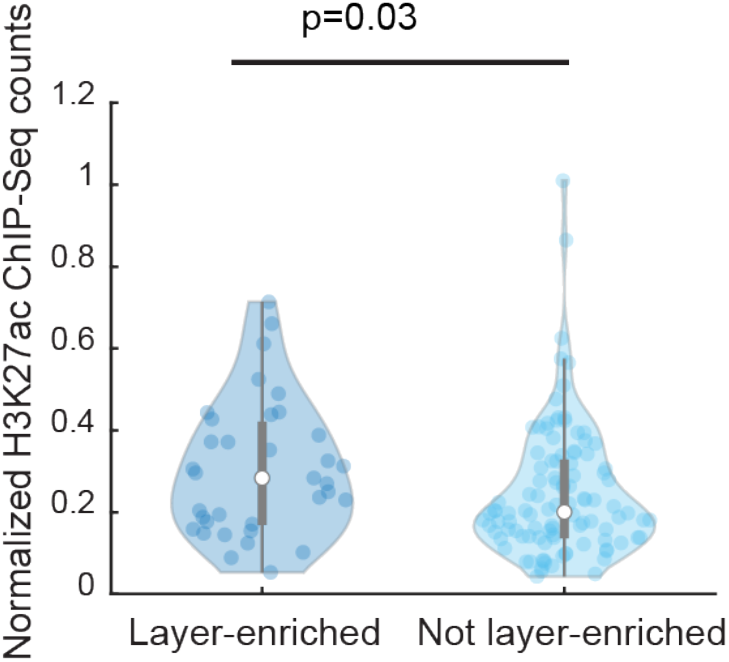
Comparison of H3K27ac ChIP-Seq signals between target loci exhibiting and not exhibiting layer-enriched H3K27ac signals. Violin plots showing the H3K27ac ChIP-Seq read number for each target H3K27ac locus for loci that showed layer-specific enrichment (left) and loci that did not show significant layer-specific enrichment (right). ChIP-Seq data are taken from (Mo et al., 2015).

**Figure S7.**
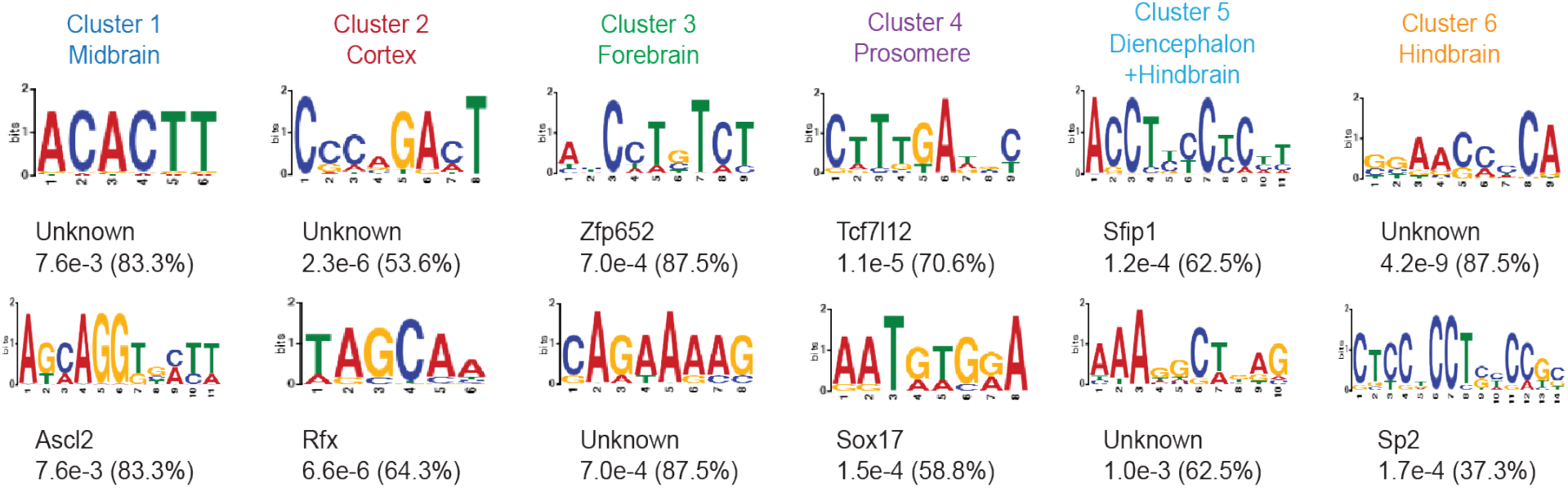
Top sequence motifs identified for putative enhancers in the six clusters shown in. Figure 5. The top two most enriched sequence motifs for each of the six clusters are shown. The p-value with the percentage of locus having the motif in bracket for each motif were listed below.

## Supplementary Table Caption

**Table S1. Sequences for all oligonucleotides used, related to all figures.** The table includes the following separate list of oligonucleotide probes:

Encoding probe library for 90 target H3K27ac loci in hTERT-RPE1 cells Encoding probe library for 52 target H3K4me3 loci in hTERT-RPE1 cells

Encoding probe library for 127 target H3K4me3 loci in mouse cortex and mouse embryonic brain Encoding probe library for 139 target H3K27ac loci in mouse cortex

Encoding probe library for 142 target H3K27ac and 5 negative control loci in mouse embryonic brain

Library amplification primers for all the above encoding probe libraries.

The readout probes

Tn5 loaders

Oligonucleotides for SON experiments

The lists for the encoding probe libraries, library amplification primers, Tn5 loaders and oligonucleotides for SON experiments include: 1) Name of each oligo, 2) Sequences of each oligo. The list for the readout probes includes 1) Bit number, 2) Readout probe name, 3) Sequence of the readout probe 4) The fluorophore for that readout probe.

**Table S2. MERFISH codebooks for the target epigenomic loci, related to all figures.** The table includes the codebook of the five sets of target loci: 90 target H3K27ac loci in hTERT-RPE1 cells, 52 target H3K4me3 loci in hTERT-RPE1 cells, 127 target H3K4me3 loci in adult mouse cortex and embryonic mouse brain, 139 target H3K27ac loci in adult mouse cortex, and 142 target H3K27ac loci and 5 negative control loci in embryonic mouse brain. Each codebook contains: 1. Loci number 2. Chromosome of the loci, 3. Start genomic positions of the loci, 4. End genomic positions of the loci, (5. Name of the corresponding gene when relevant), followed by barcodes of the individual loci as a series of ‘0’ (off bits) and ‘1’ (on bits). The corresponding readout probe for each bit is labeled on top.

